# Deep mutational scanning of the human insulin receptor ectodomain to inform precision therapy for insulin resistance

**DOI:** 10.1101/2024.09.07.611782

**Authors:** Vahid Aslanzadeh, Gemma V. Brierley, Rupa Kumar, Hasan Çubuk, Corinne Vigouroux, Kenneth A. Matreyek, Grzegorz Kudla, Robert K. Semple

**Author notes:** Correspondence: Prof. Robert K. Semple [Lead Contact], X: @SempleLab or Prof. Grzegorz Kudla.

## Abstract

The insulin receptor (INSR) entrains tissue growth and metabolism to nutritional conditions. Complete loss of function in humans leads to extreme insulin resistance and infantile mortality, while loss of 80-90% function permits longevity of decades. Even low-level activation of severely compromised receptors, for example by anti-receptor monoclonal antibodies, thus offers the potential for decisive clinical benefit. A barrier to genetic diagnosis and translational research is the increasing identification of *INSR* variants of uncertain significance. We employed saturation mutagenesis coupled to multidimensional flow-based assays to stratify approximately 14,000 INSR extracellular domain missense variants by cell surface expression, insulin binding, and insulin- or monoclonal antibody-stimulated signaling. The resulting function scores correlate strongly with clinical syndromes, offer insights into dynamics of insulin binding, and reveal novel potential gain-of-function variants. This INSR sequence-function map has high biochemical, diagnostic and translational utility, aiding rapid identification of variants amenable to activation by non-canonical INSR agonists.

---

Insulin is released in response to food ingestion, orchestrating co-ordinated physiological responses promoting energy storage and growth, and is essential for human life. Its effects are mediated by binding to a ubiquitously expressed dimeric transmembrane receptor tyrosine kinase. Insulin binding leads to receptor autophosphorylation, which triggers a network of biochemical responses including activation of phosphoinositide 3-kinase, and phosphorylation and activation of the downstream AKT1 serine-threonine kinases^1^

Insulin binds to its receptor with complex kinetics ^2^ There are two pairs of binding sites in the receptor dimer, and both positive and negative co-operativity are apparent in sequential binding of insulin molecules. Major insights into the structural basis of these kinetics, and of transduction of insulin binding into trans-autophosphorylation of the intracellular domain of the receptor, have been gained through application of X-ray crystallography and, latterly, cryoEM^3,4^. Important structure-function questions remain, however, particularly relating to details of the conformation changes induced by insulin binding that activate intracellular trans-autophosphorylation.

Since the first descriptions in 1988^5,6^ more than 150 disease-causing mutations in the *INSR* gene, encoding the insulin receptor, have been reported, and the range of associated disease severity is well delineated^7,8^. In brief, biallelic mutations usually cause extreme autosomal recessive insulin resistance (IR) that is apparent neonatally, while heterozygous, dominant negative intracellular mutations usually cause severe IR presenting around puberty^7^. *INSR*-related IR syndromes are often named after their describers. Near complete loss of receptor function causes Donohue syndrome, leading to mortality in early months or years of life, while in Rabson Mendenhall syndrome, another recessive syndrome, survival to adulthood is possible thanks to residual receptor function^7^. Later onset dominant INSR-related IR, often called “Type A” insulin resistance, features up to 25% receptor function and is usually amenable to management with current therapies^7^. There is no discernible clinical impact of 50% loss of receptor function.

Mutations in the extracellular domain of the receptor can lead to a diverse range of molecular phenotypes, including changes of receptor abundance, localization, insulin binding, signaling, and/or receptor recycling. No single biochemical assay captures all these effects; although several tens of INSR mutations have been studied functionally over the past 4 decades, reported studies used a variety of in vitro and cellular approaches, and it is not established which assay best predicts clinical outcomes. As a result, accurate prognostication based on biochemical measurements of residual receptor function remains challenging. This is an acute problem as diagnostic testing commonly reveals novel “variants of unknown significance” (VUS), often in people of ethnicities poorly represented in current large population datasets.

The steep relationship between INSR function and clinical outcome at the lower end of the range implies that even modest improvement in function could confer decisive clinical benefit in extreme recessive IR. A promising approach to augmenting signaling by severely deleterious INSR variants is to deploy agonists with a mode of action distinct to that of insulin. Bivalent antibodies against the extracellular domain of the receptor can activate wild-type receptor in cells^9^ and in vivo^10,11^. Crucially, different antibodies have been shown also to activate severely dysfunctional mutant receptors^12–14^, exerting insulin mimetic metabolic actions in cells^14,15^ and in a humanised mouse model^14,16^ However, although this approach has been validated for a small number of well-studied INSR mutations, the full repertoire of mutations activatable in this way is unknown.

We now sought to address the diagnostic challenges posed by VUS, to improve prognostication based on more detailed understanding of INSR genotype-function relationships, and to stratify loss-of-function mutations for novel agonist therapy. To do this we developed a multidimensional, massively Multiplexed Assay of Variant Effect (MAVE). Specifically, we employed a plasmid-based approach to saturation mutagenesis of the extracellular, ligand-binding domain of the human insulin receptor, coupled to massively parallel orthogonal assays of variant cell surface expression, insulin and antibody binding, and insulin and antibody-stimulated signaling.

## Results

### INSR mutation library creation and validation

We chose to focus on the extracellular portion of the B isoform of the human insulin receptor. The B isoform includes the 36-nucleotide exon 11, and is regarded as the more “metabolic” of the INSR splice variants^17^. Around two thirds of the full length receptor is extracellular (**Figure S1A**), and it is in this insulin-exposed region that disease-related mutations that compromise insulin binding lie. These are the mutations which are potentially activatable by novel receptor ligands such as bivalent antibodies.

We adopted a plasmid-based approach to mutate residues 28-955, from the first codon of the mature receptor to the transmembrane domain. We subcloned the *INSR* gene into an entry plasmid before restriction enzyme-based incorporation of 30mer barcode sequences upstream from the *INSR* open reading frame, with an interposed *attB* recombination site to permit efficient Bxb1- mediated integration into a landing pad in the genome^18^. Mutations were introduced into this library by nicking mutagenesis based on a published method^19^ to yield a mutagenised, barcoded library. Long read PacBio sequencing was used to verify the library and to establish pairing of barcodes and mutations (**Figures 1A,B**; **S1B-D**). We observed 15,996 single missense and stop codon variants out of 18,560 possible (86% coverage), tagged by 80,956 unique barcodes. 81% of mutations associated with two or more barcodes (**Figure S1C**). 35% of the barcodes aligned with unmutated INSR or synonymous mutations only, and were taken as wild-type background in downstream analyses (**Figure S1E, Tables S1,S2**).

**Figure 1:**
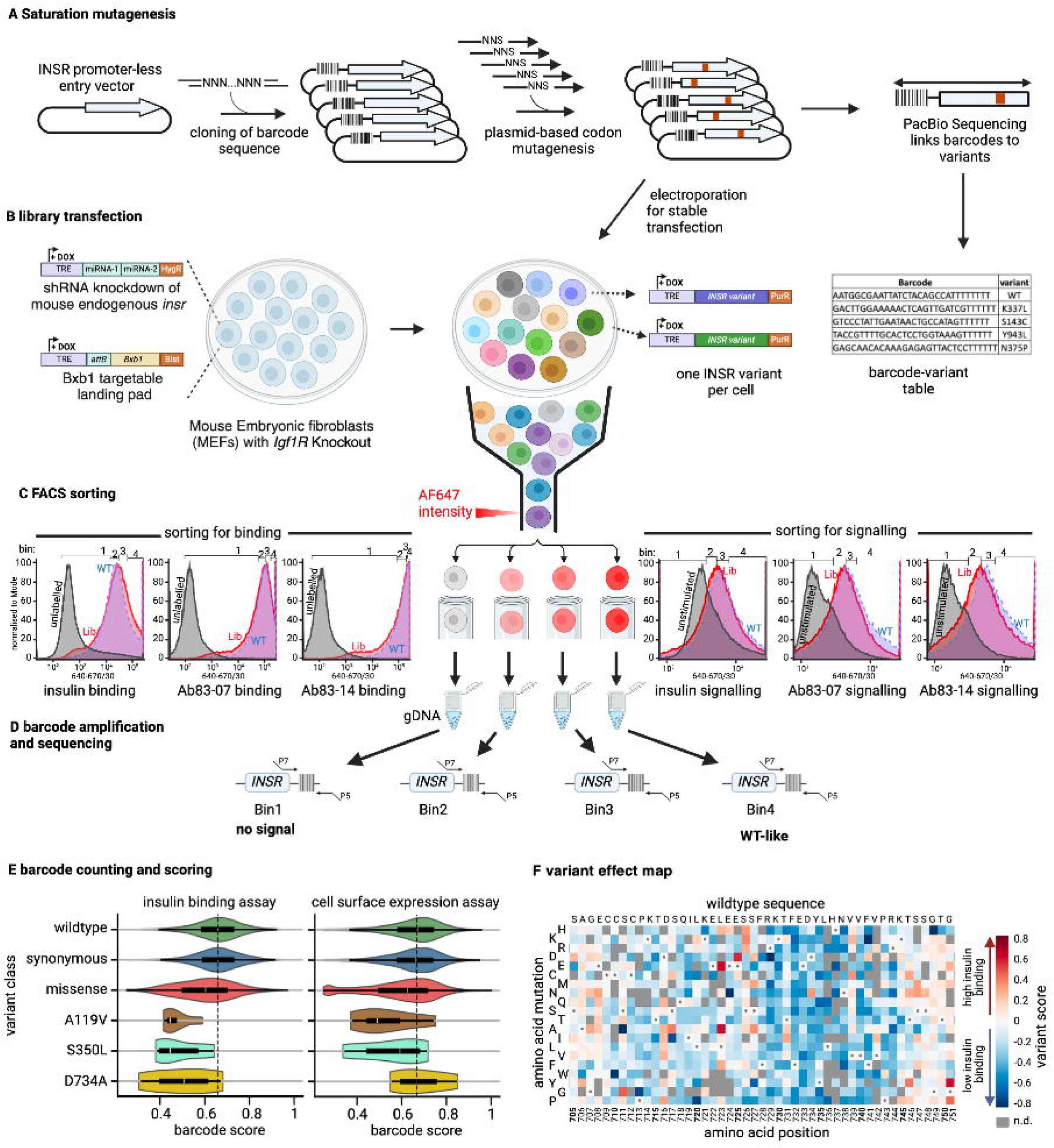
Overview of workflow for massively parallel analysis of INSR variant effect. **A**, Schematic of saturation mutagenesis of the INSR ectodomain, including barcoding of an entry vector with 30 random nucleotides, PCR-based INSR ectodomain mutagenesis to generate the barcode-variant library, and long read DNA sequencing to phase barcodes and mutations. **B**, Transfection of the barcoded library into Igf1r knockout Mouse Embryo Fibroblasts (MEFs) with doxycycline-inducible INSR knockdown and a Bxb1-targetable landing pad to generate a large library of cells, each with a uniquely barcoded, conditionally expressible INSR variant. **C**, Sorting of the cellular variant library into four bins based on binding assays (left) or signaling assays (right). Bin thresholds are shown above the plots, with WT (blue), variant libraries (red), and control (grey, no transfected MEFs) overlaid. At least 13 million cells were sorted into each bin. **D**, Extraction of gDNA from each bin and amplification of barcodes with primers linked to Illumina p5 and p7 sequences. **E**, Violin plots representing distribution of insulin binding (left panel) and cell surface expression (right panel) barcode scores for WT, synonymous, missense, and three known pathogenic variants. **F**, A detail of a sequence-function map representing variant scores for insulin binding for amino acids 705-751. Cell colour indicates the score for a single amino acid change. Variants are arrayed by position (column) and variant amino acid (row). Positive scores (in red) and negative scores (in blue) indicate better and worse function than wild-type in the assay, respectively. Grey cells are missing data and white cells with a dot in centre indicate the wild type residue at that position. See also **Figures S1** and **S2**

Study of INSR variant function requires a cellular background free of both endogenous insulin receptor and the highly homologous insulin-like growth factor 1 (IGF1) receptor, which can respond to high insulin concentrations. We thus turned to a mouse embryo fibroblast model in which *Igf1r* knockout^20^ is augmented by doxycycline-inducible, shRNA-mediated knockdown of the endogenous *Insr* gene^14^. We chose conditional *Insr* knockdown as *Insr* knockout on an *Igf1r* null background in this cell line reproducibly yielded cellular atypia and compromised viability in our hands. Into this cell line we lentivirally introduced a single copy of a previously described landing pad allowing efficient *Bxb1*-mediated integration of transfected plasmid^18^. Finally, the mutagenised, barcoded human *INSR* plasmid library was introduced to cells by electroporation, before antibiotic selection of cells with integrated plasmid (**Figure 1B**).

### Design and validation of INSR functional assays

Having generated a large, conditionally expressible library of INSR ectodomain variants, we next aimed to determine which mutations are expressed at the cell surface, which bind insulin, and which mediate insulin- or antibody-induced metabolic signaling. To this end we validated flow sorting assays capable of handling tens of thousands of variants in millions of cells in parallel (**Figure 1C**). To measure cell surface expression we used binding of two well characterised human INSR-specific monoclonal antibodies (mAbs 83-7 and 83-14) with distinct receptor epitopes^21^. We have previously shown that both these antibodies bind mutated receptors, eliciting signaling even by some mutant receptors with absent or severely attenuated responsiveness to insulin^14^. To assess insulin binding we used recombinant human insulin. All ligands were labelled with the fluorescent dye AlexaFluor 647 (AF647). Finally, to assess signaling induced by insulin or antibodies we permeabilised cells and probed with an antibody against phosphorylated AKT, a key mediator of insulin’s metabolic actions^22^ (**Figure 1C**).

In each assay cells were flow-sorted into four bins according to fluorescence intensity. From every bin DNA was extracted, barcodes were amplified by PCR, and high-throughput sequencing was used to quantify variant frequency per bin (**Figure 1D**). Next, a weighted average was calculated for each barcode based on its distribution across the bins and this was used to derive a score for each variant in every replicate experiment. Barcode score distributions for both insulin binding and cell surface expression were superimposable for wild-type receptors and synonymous variants but skewed towards lower scores for all missense variants. Both scores were lower for previously studied loss of function variants A119V^23^, and S350L^14,16,24,25^, while only insulin binding was lower for D734A, in keeping with prior studies^14,16,26^ (**Figure 1E**). False discovery rates were computed by a bootstrapping approach using all barcode scores per variant across replicate sorts of the same cell library. Replicates were finally averaged and displayed in array form in variant effect maps. (**Figure 1F**, **Figure S2A, Tables S2,S3**).

Pilot experiments confirmed that flow-based assays do report on relevant behaviours of receptor variants (**Figure S1F**). To assess between-assay variability, we conducted two sorting experiments for insulin binding, five for insulin-activated signaling, and two for binding and three for signaling of each antibody. Variant-effect maps and variant function scores were consistent across replicates (**Figure S2A,B**), and PCA analysis revealed discernible clusters corresponding to distinct experimental conditions (**Figure S2C**). Means of the replicate scores were thus used in subsequent analyses.

### Variants reducing cell surface expression and/or insulin binding

Of 15,996 variants present in the input plasmid library, 13,638 were detected, on average, across the bins in all sortings. This represented 79% of all possible missense variants and 88% of the repertoire of missense mutations in the barcoded plasmid library (**Table S1**). Overall visualisation of binding scores for insulin and antibodies showed a highly non random pattern (**Figure 2A**). For most regions of the ectodomain binding was broadly consistent for antibodies and insulin, in keeping with loss of receptor expression (**Figure S3A**). Particularly clear examples were the series of cysteines involved in intramolecular disulphide bonds within the cysteine-rich domain (**Figure S3B**). These residues were obvious as columns of blue in the variant effect maps, indicative of residues where any change from the native residue was highly deleterious to cell surface expression of the receptor (**Figure 2A**), and are illustrated in the cryoEM structure of the 4-insulin-bound “T” conformation of the receptor in **Figure S3C**.

**Figure 2:**
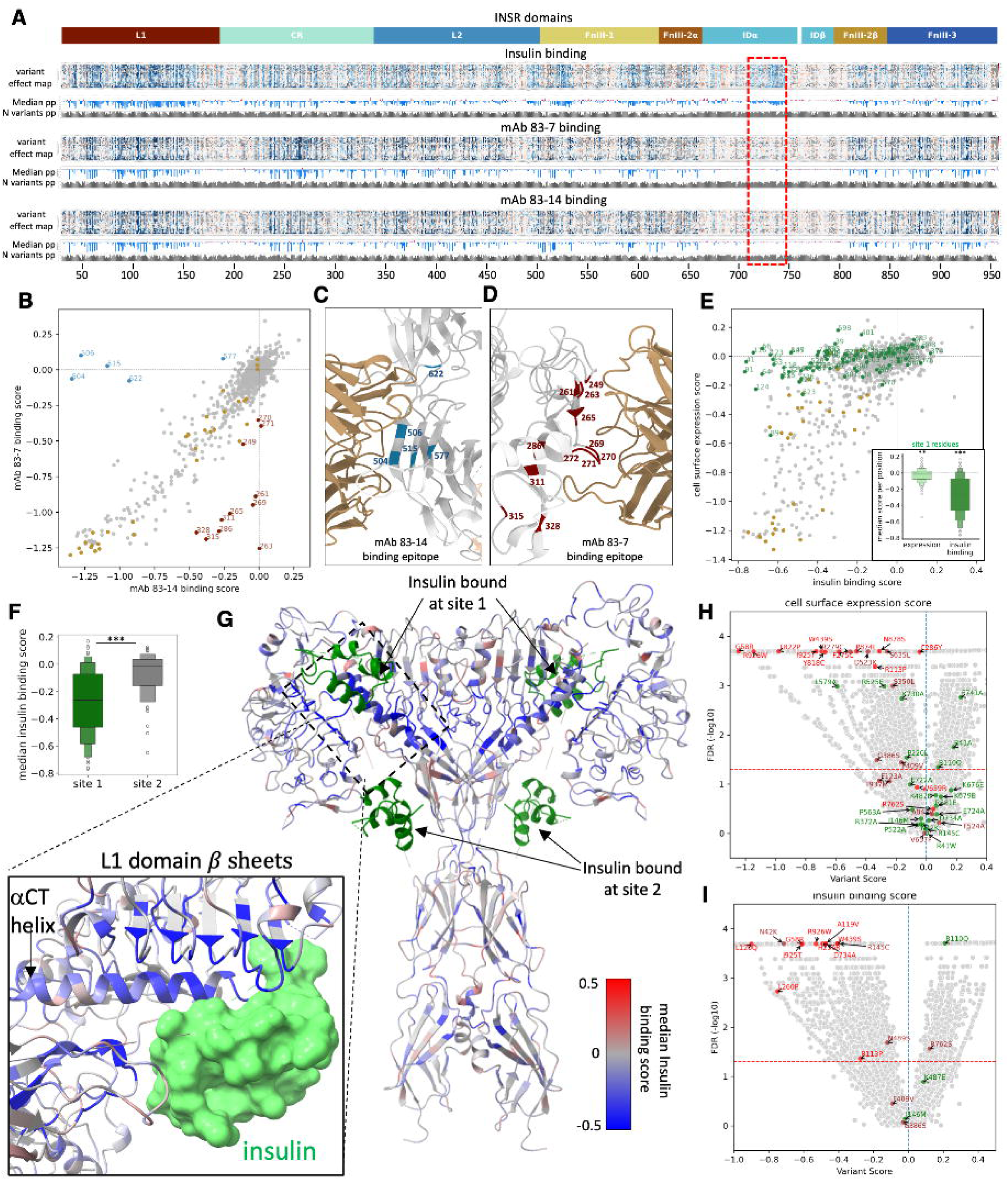
Effects of mutations on cell surface expression and insulin binding. **A**, Heatmap of variant scores for insulin, mAb 83-7 and mAb 83-14 binding, referenced to INSR extracellular domain architecture. Unscored variants are coloured grey. ‘median pp’ indicates median variant scores per position and ‘N variants pp’ indicates number of variants scored per position. The rectangular dashed red box demarcates the αCT region (716-746). **B**, Scatter plot of median scores per position for mAb 83-7 binding with dark-brown coloured outliers and mAb 83-14 binding with light-blue coloured outliers. Cysteine residues involved in disulphide bond formation are coloured golden brown for reference. **C, D**, Detail of INSR structure (from PDB 4ZXB) showing mAb 83-7 and mAb 83- 14 epitopes respectively, with outlier residues from **B** highlighted. **E**, Scatter plot of median scores for insulin binding and cell surface expression, calculated as the mean of mAb binding scores. Residues involved in insulin binding site 1 are shown in green. Cysteine residues involved in disulphide bond formation are coloured golden brown for reference. The inset boxplot shows expression and insulin binding scores for site 1 residues. Wilcoxon signed-rank test was utilized to determine if insulin binding and expression scores for site 1 residues differ from wildtype, asterisks on the box plots indicate significance. *, ** and *** denote p-values less than 0.05, 0.01 and 0.005 respectively. **F**, median insulin signaling score distribution for site 1 and site 2 residues, compared by Mann Whitney U test. **G**, 4 insulin-bound INSR structure (PDB: 6PXV) coloured by median insulin binding score after excluding residues with low expression scores (**Figure S3E**). Insulin molecules are green. **H**,**I**, Volcano plots of cell surface expression score or insulin binding score respectively (x axis) against -log10 FDR (y axis). Annotated points are functionally studied variants coloured red, brown or green, respectively, denoting degree of functional impairment in prior assays Red = severely impaired; Green = little or no reported impairment; Brown = intermediate impairment (**Table S4**). Dashed horizontal line is FDR of 0.05 (-log10 (0.05)). See also **Figures S3** and **S4.**

As expected, there was a strong correlation between results derived from use of the two monoclonal antibodies, whether scored for each variant (**Figure S3D**) or averaged for all variants at each residue (**Figure 2B**). Several striking outliers were observed in antibody correlation plots, however. As both the antibodies studied were included in previous crystal structures of the INSR ectodomain^21^ we were able to assess whether such outlying amino acids were implicated in antibody-binding epitopes. This did prove to be the case. All 5 outlying residues whose mutation selectively disfavours mAb 83-14 map to the crystallographically determined mAb 83-14 epitope (**Figure 2C**), while all 11 residues whose mutation selectively disfavours mAb 83-7 binding mapped to the mAb 83-7 epitope (**Figure 2D**). These observations demonstrate the utility of the binding assay for identifying epitope residues making important thermodynamic contributions to antibody binding. For subsequent analyses, the means of the scores for the two antibodies were taken as the (cell surface) ”expression score”.

Although binding patterns for antibodies and insulin were broadly similar, some important differences were apparent even in the high level view. Most prominently, in the insert domain, insulin binding was strongly disfavoured by mutations, seen as a blue blush in the top trace in **Figure 2A** (boxed), while antibody binding in the same region was relatively unperturbed, consistent with preserved cell surface expression. This region, experimentally identified as being important for insulin binding, corresponds to the so called α−C-terminal (αCT) (residues 716–746), an α-helix forming a key part of insulin binding site 1, which is in direct contact with bound insulin. Site 1 is formed by the L1 domain of one INSR monomer and α-CT and FnIII-1 loop of the other monomer^3,4^.

We next extended this observation by comparing antibody and insulin binding scores for all 80 residues implicated in prior structural studies in engaging in insulin binding in site 1^27–33^, comparing them to results for cysteines in the cysteine-rich (CR) domain (**Figure 2E**). In each case medians of all available amino acid substitutions were computed for each residue. Mutation of site 1 residues led to a very small but significant reduction of cell surface INSR expression compared to all other missense variants, but to much more substantial loss of insulin binding (**Figure 2E**, **inset)**. This was in contrast to CR cysteine mutations, which nearly all reduced both INSR expression and insulin binding (**Figure S3B**). Mutation of amino acids forming insulin binding site 2 had no effect on receptor expression and only very mildly reduced insulin binding on average, although outlying variants that severely compromised one or both were seen (**Figures 2F**, **S3E**).

Next we used a structurally agnostic approach to identify residues important for insulin binding but not receptor expression. Colouring the INSR structure based on residues’ median insulin-binding scores without adjusting for the impact of mutations on expression precludes identification of mutations that selectively influence insulin binding. We corrected this by excluding any variants with marked loss of cell surface expression (**Figure S3F**), then colouring the insulin-bound INSR structure according to remaining residues’ median insulin-binding scores (**Figure 2G**). This revealed dense clustering of the lowest binding scores in apposition to insulin, notably in the αCT, and L1 elements of insulin-binding Site 1. Clusters of residues whose mutation selectively affected binding were also identified in interdomain hinge regions of the receptor, but residues around insulin-binding Site 2 did not stand out strongly.

We next assessed individual missense variants. This identified 4,840 variants with significantly reduced expression scores, 1,458 with significantly increased expression scores (**Figure 2H**), 4,625 with significantly reduced insulin binding scores and 986 with significantly and consistently increased binding scores (**Figure 2I**). To evaluate our findings against prior reports, we collated results of all previous studies reporting functional consequences of individual INSR variants, whether disease-associated or studied as part of structure-function investigations. Variable experimental approaches used over 36 years, and differing degrees of rigour in execution and reporting, with many results semi-quantitative only, posed a challenge to curation. Pragmatically we stratified variants studied into 3 groups for each functional property: those reported to confer severe (<10% wild-type) or complete loss of function (LoF), those with evidence of lesser LoF, and those with unimpaired function. We also adjudicated qualitatively on the likely confidence of findings based on the methods used (**Table S4**).

Comparison of expression and insulin binding data from our massively parallel assays to these prior experiments revealed strong concordance. For cell surface expression, 11 of the 12 reported severe LoF variants and 17 of the 22 variants with any reported degree of LoF were called as LoF with high confidence in the high throughput data. The only reported severe LoF variant discordantly classified in the high throughput data was W659R. W659R was reported in a patient with Donohue syndrome in association with a dominant negative tyrosine kinase domain variant V1054M^34^, meaning the in vivo effect of W659R alone is difficult to quantify. Moreover prior experimental evidence for LoF was a single immunoblot after transient overexpression^35^. No significant difference was found for insulin binding or signaling in the current study, despite at least 6 barcodes representing the variant in all experiments. The median binding score for the 19 total substitutions at this position were available was only -0.06 (interquartile range -0.15-0.07).

Conversely, 4 of 24 variants reported previously to be expressed normally were called as showing loss of expression in this study, 1 (P220L) with only marginal significance. Evidence for preserved expression of the other 3 variants came from transient overexpression and immunoblotting in a single study, in which maximal insulin signaling was reported to be moderately reduced for all 3^28^. All 3 also showed reduced insulin binding in the high throughput assay. For insulin binding, 11 out of 13 reportedly severe LoF variants and 14 out of all 18 LoF variants were called as LoF with confidence. One reported severe LoF variant had a low confidence LoF score, while one - R762S - showed a discrepancy, being called as a gain of binding variant. R762 lies within the tetrabasic furin cleavage motif, and R762S was reported previously to be expressed at the cell surface but to show severely impaired insulin binding due to failed proreceptor cleavage^6^. Importantly, this defect was correctable by light trypsinisation. As trypsin was used in the preparation of adherent cells in this study for flow sorting, it is likely that this remedied defective proreceptor cleavage, restoring insulin binding. Whether other INSR variants are similarly affected is unknown.

We also compared our findings to the alanine-scanning mutagenesis studies which were influential in identifying ectodomain residues important for insulin binding, largely before receptor structures were reported ^29,30,32^. Correlation of insulin binding scores from this study with ratios of mutant to wild-type insulin dissociation constants from alanine scanning studies are shown in **Figure S4**. 21 of the 23 reportedly severe LoF variants in previous studies were also found to confer loss of binding in the current study, 15 with a FDR <0.05 (**Figure S4A**). When function scores were averaged across all substitutions at the residues concerned, results aligned with alanine scanning for all variants, however (**Figure S4B,C**). Of the 28 alanine variants called confidently as LoF in the current study, only 15 were also deemed LoF in prior alanine scanning (**Figure S4A**). This may in part relate to greater sensitivity of the present assay, but also reflects true differences in binding based on experimental paradigm: prior studies used soluble INSR ectodomains, and although these constructs do bind insulin, they do so with low affinity, and without the co-operativity characteristic of full length receptor binding^36^. Some mutations, such as D734A, have been clearly established to confer loss of binding on full length receptor, in agreement with this study^26^, but not on receptor ectodomain alone^30^, in keeping with prior studies.

#### Variants increasing insulin binding

Mutation of a small number of residues increased maximal insulin binding for most or all substitutions tested. For many of these this correlated with increased cell surface expression, but mutations of some residues increased binding without increasing expression, implying an important role for the native amino acid in regulation of binding (**Figure 3A**). By far the most striking amongst these was G333, which markedly increased maximal binding in the face of mildly reduced expression. 16 of the 17 individual substitutions assayed at position 333 increased insulin binding, despite 14 of them decreasing expression, with G333Q showing one of the strongest positive effects.

**Figure 3:**
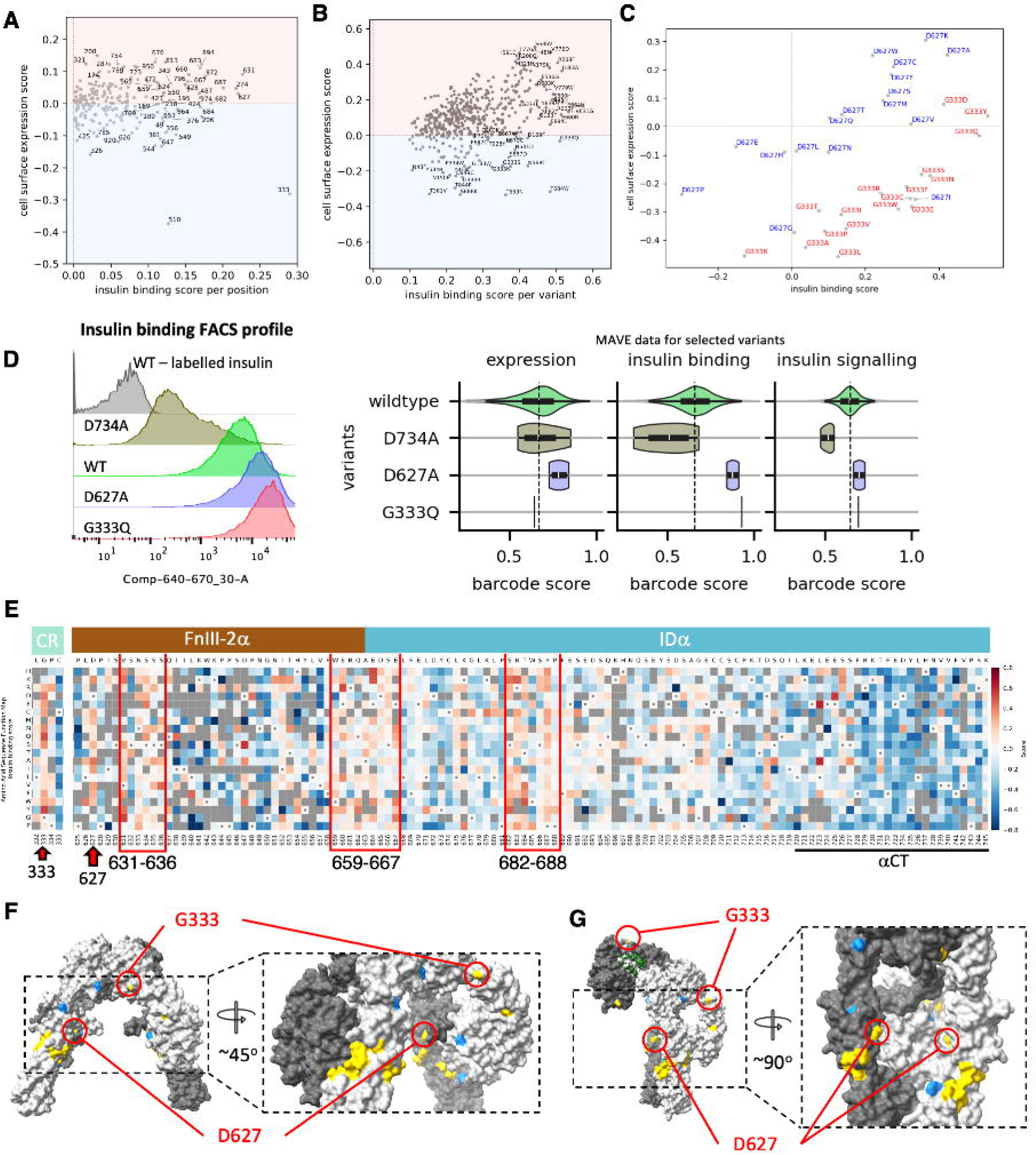
Variants Increasing Insulin Binding. **A**, Detail of scatter plot of median of mean antibody binding scores per position (“expression scores”) against insulin binding scores, focused only on gain of insulin binding. **B**, Scatter plot of gain-of-insulin-binding scores with FDR < 0.01 against expression scores for individual variants. **C**, G333 and D627 variant insulin binding vs expression scatter plot. **D**, FACS profile from dedicated replication experiment assessing D627A and G333Q, with D734A used as a control, shown next to barcode score distributions for wildtype, D734A, D627A, and G333G for expression, insulin binding, and signaling scores. G333Q is represented by a single vertical line as it has only one barcode. **E**, Heatmap with highlighted gain-of-insulin-binding hotspots, visually ascertained. **F**,**G** INSR residues identified to result in gain of function upon substitution mapped to the cryo-EM structures of the various insulin-bound states. Monomers are coloured different shades of grey. Insulin is represented by green ribbons. Yellow depicts residues where multiple substitutions increase insulin binding. Blue depicts residues where a unique substation increases binding. Some residues highlighted may be involved in interactions with parts of the receptor not resolved in available structures. **F,** Unbound, INSR B-isoform (PDB: 8U4B). The highlighted section is rotated counterclockwise 45° to the full view of the receptor. **G,** Single insulin bound (green ribbon) INSR isoform A (PDB: 7STI). PDB structure 7STI is the A isoform of the Mus musculus orthologue, but for consistency residues highlighted are numbered according to the human B isoform pro-receptor with the corresponding mouse residue in brackets. The highlighted section is rotated counterclockwise 90°. See also **Figure S5**.

(**Figure 3B,C**). G333 lies in the middle of the CR:L2 hinge region. Although it is in an exposed loop in the fully insulin-bound receptor conformation, this region undergoes considerable conformational change between the apo and insulin-bound receptor and substitution of glycine for bulkier amino acids may plausibly alter the energetics of this conformational change (**Figures 3F,G** ;). **S** A**5**lso demonstrating frequent gain of binding upon substitution were positions 274, 627, 631, 633-636, 659-667, 682-684 and 686-688 (**Figures 3E-G**; **S5**). D627, one of the residues whose mutation conferred the strongest average gain of binding, lies in the FnIII-2a domain (**Figures 3F,G**; **S5**). 10 of the 17 substitutions tested for D627 increased both binding and expression, with substitution for alanine having the greatest effect (**Figure 3C**). This is in keeping with a previous alanine scanning mutagenesis study of the INSR ectodomain^37^. To verify apparent gain of insulin binding, we selected G333Q and D627A for further study. Both mutations were generated as well as D734A as a negative control known to impair binding, and all variants were studied individually by the same flow cytometry-based assays. These confirmed findings from the multiplexed assay, with markedly increased insulin binding for G333Q and D627A and reduced binding for D734A, and increased cell surface expression for D627A only (**Figure 3D**).

Turning to individual missense variants, once again many which increased maximal insulin binding also increased cell surface expression, likely explaining the increased binding. A subset increased insulin binding despite little or no increase in receptor expression, however (**Figure 3B**). Some of these mutations occurred at the amino acids previously identified by plotting averaged scores per position (**Figure 3A**), and some further amino acids such as G183 were revealed for which several substitutions increase maximal binding (**Figure 3B**). Some variants stood out from other substitutions at the same amino acid, however, implying neomorphic gain of function unique to that substitution (e.g. L1 domain: D169F; FnIII-1: N541D, N568C; FnIII-2A: T653N, K679C, T684W; **Figure 3B**). Mapping native amino acids at these positions onto available INSR structures reveals that they cluster at sites of interdomain interaction that are highly dynamic on sequential insulin binding. This clustering is particularly prominent in the region where the L1’ packs against FnIII-2a in the unbound conformation, and the region where the ID domain threads through the interior of the receptor, although the latter is unresolved in available structures. It is plausible that many of the variants increasing maximal insulin binding alter the thermodynamics of insulin-induced receptor conformational change, but this will require future study (**Figures 3F,G**; **S5**).

#### Effects of INSR mutations on insulin signaling

To assess insulin signaling, we used a fluorescently labelled antibody against phospho-Akt (Ser473/4) to quantify insulin-induced Akt phosphorylation. As a single high concentration of insulin was used (100 nM) this assay effectively determined the maximal level of insulin-induced phosphorylation only (E_max_). While the insulin and antibody binding assays showed a >100-fold dynamic range (fluorescence ratio of insulin-stimulated vs unstimulated cells), the dynamic range of the signaling assay was less than 5-fold. The signal-to-noise ratio of signaling scores was therefore lower; to compensate for this, we conducted the assay in 5 replicates and averaged the results. High level results are illustrated in **Figure 4A**, with insulin binding results as a comparator. Once again, results were non random, and insulin-induced Akt phosphorylation visually corresponding to insulin binding (**Figure 4A**). Mutations in insulin binding site 1 residues had less effect on signaling than on binding (**Figure 4B**), in keeping with the “spare receptor” concept, whereby maximal signaling is seen at relatively low levels of receptor occupancy^38^ . It follows from this that considerable loss of insulin binding may be seen before peak insulin signaling is compromised. A minor contribution from variants that activate basal signaling remains possible, though no such variants have been described to date in the ectodomain of the INSR. As before, mutation of site 2 residues only mildly reduced binding and signaling in group-based analysis, although scattered outlying amino acids were seen whose mutation had much more deleterious effects on both binding and signaling (**Figure 4C** and inset). These effects were smaller than for Site 1 mutations (**Figure 4B-D**). Mapping signaling scores of well expressed residues onto the 4 insulin-bound receptor structure, as before, the largest effect on signaling was observed for residues clustered around Site 1, though a deleterious effect was also seen for many residues in the “legs” of the receptor which are involved in receptor compaction in the unliganded state (**Figure 4E**).

**Figure 4:**
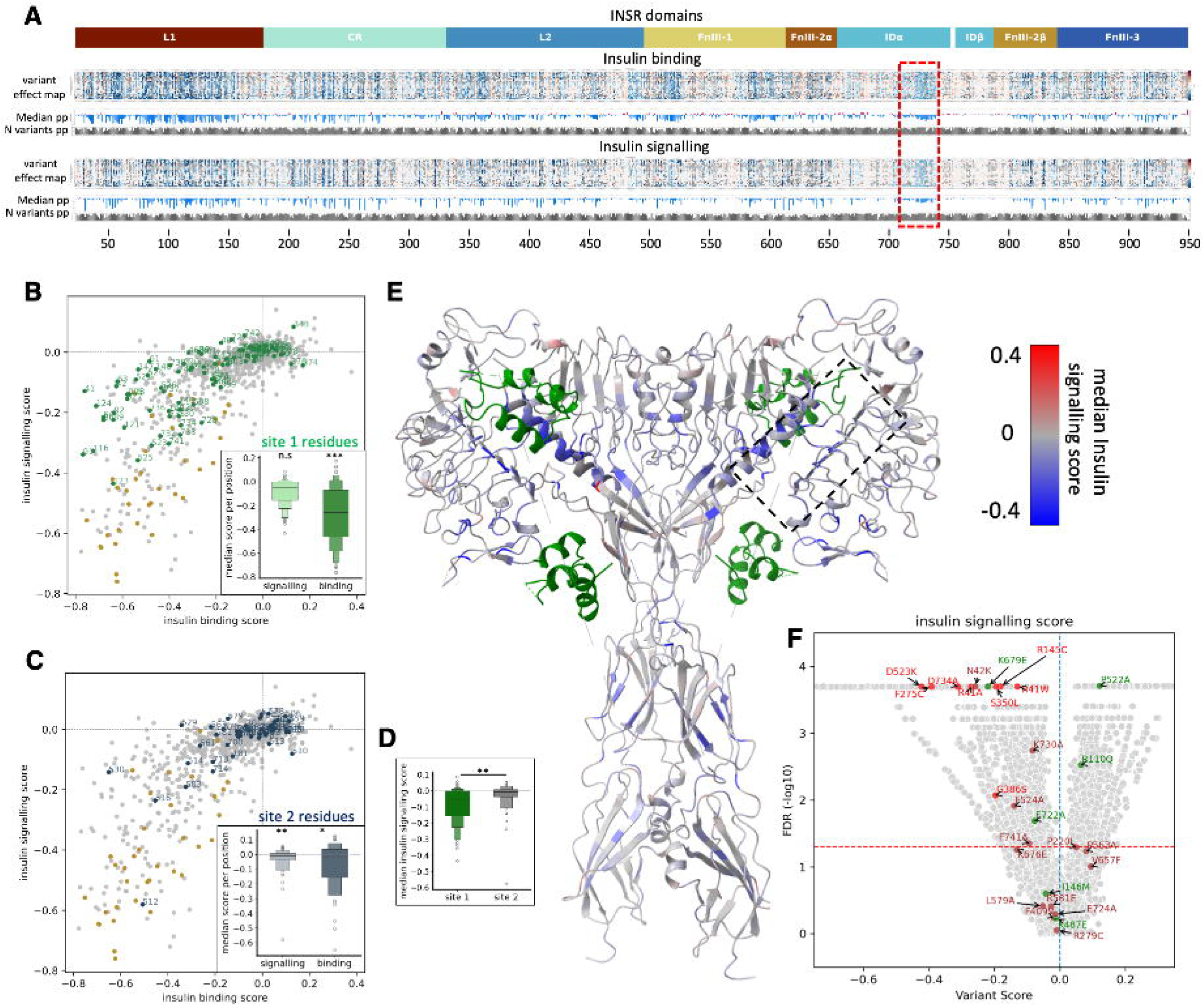
Effects of mutations on maximal INSR signaling. **A**, Heatmap of variant scores for insulin, mAb 83-7 and mAb 83-14 binding, referenced to domain architecture. Unscored variants are coloured grey. ‘median pp’ indicates median variant scores per position and ‘N variants pp’ number of variants scored per position. The rectangular dashed red box on the heatmaps demarcates the αCT region. **B**, Scatter plot of median scores for insulin binding against median signaling scores with site 1 residues highlighted in green. The inset boxplot shows median insulin binding and signaling scores for site 1 residues. Wilcoxon signed-rank test was utilized to determine if insulin signaling and binding for site 1 residues differ from wildtype. Asterisks indicate significance. *, ** and *** denotes p-values less than 0.05, 0.01 and 0.005 respectively. **C**, Scatter plot of median scores for insulin binding against median signaling scores, with dark-blue-highlighted site 2 residues. The inset boxplot displays median insulin binding and signaling scores for site 2 residues with statistical comparison as for site 1. **D**, Median insulin signaling score distribution for site 1 and site 2 residues, compared directly using the Mann Whitney U test. *, ** and *** denote p-values less than 0.05, 0.01 and 0.005 respectively. **E**, 4 insulin-bound INSR structure (PDB: 6PXV), coloured by median insulin signaling scores after excluding residues that markedly lower cell surface expression according to **Figure S3E**. Insulin molecules are green. **F**, Volcano plots of insulin signaling scores (x axis) against -log10 FDR (y axis). Annotated points are functionally studied variants coloured red, brown or green, respectively, denoting degree of functional impairment in prior assays. Red = severely impaired; Green = little or no reported impairment; Brown = intermediate impairment (**Table S4**). Dashed horizontal line denotes FDR of 0.05. See also **Figures S6** and **S7**

Turning to individual missense variants, we identified 5,327 variants with significantly reduced peak insulin-induced signaling, and 1,500 variants with significantly increased signaling (**Figure 4A,F**; **Tables S2,S3**). All 8 variants found in previous studies to show severe loss of signaling and 12 of 21 reported LoF variants were called concordantly in the high throughput assay. Nine of the remaining variants had only lower confidence signaling scores (**Figure 4F**).

The signaling assay used, solely determining maximal signaling, was not suitable for detection of variants that enhance signaling in the normal dynamic range of insulin concentrations, however a small number of variants did appear to be associated with an increase in maximal signaling (**Figure S6**). Mutation of D627, including D627A, previously verified to increase insulin binding, increased maximal signaling, while mutations of G333, including G333Q, showed only a mild increase of borderline significance (**Figure S6A-B**). Threonine 221 emerged as a residue of interest based on the average positional score (**Figure S6A**), but its average was based on only 4 variant scores in the high throughput assay. A series of other individual variants warrant future assessment in detailed signaling studies based on increased maximal signaling without increased expression. Several of the affected native residues form part of insulin binding site 1 (L2 domain: E343I, N376W; FnIII-1: N568C, P576C), others are closely involved in interdomain interactions that remodel dynamically on insulin binding (e.g. CR domain: C293F; FnIII-2A: R661W; FnIII-2B: Y839V; **Figure S6C-G**), while some are not resolved in available structures (e.g. FnIII-3: F862Y).

Comparing results for cell surface expression, insulin binding and peak insulin signaling, limiting analysis to the 5903 variants for which scores were generated for all 3 assays, and using scores for well studied variants to define loss of function cut offs (**Figure S7A-C**), we found that 74% of loss of expression variants also showed impaired insulin binding, while 51% of all loss of binding variants showed no loss of expression (**Figure S7D,E**). Only 60% of the loss of binding variants also showed loss of maximal signaling, reflecting both the presence of many spare receptors, and also the limited signaling assay employed. 23% of all the variants showing impaired signaling were both normally expressed and exhibited normal insulin binding.

#### Diagnostic performance of empirical functional results

We next asked whether functional scores of INSR variants can be used to support genetic diagnosis. Variant-level analysis broadly agreed with annotations in the ClinVar database, with known benign variants scoring significantly higher than known pathogenic variants (**Figure 5A**). However the picture was clouded by conflicting annotations for 20% of ectodomain INSR missense variants in ClinVar. Most variants in ClinVar were uniquely assigned as VUS. Of these 83 variants, 31 had significant loss of function in at least one assay, 19 in 2 assays, and 6 in all 3 assays, including variants altering highly conserved cysteines involved in disulphide bonds, for which nearly all mutations conferred strong loss of function (**Figure S8A,B**). These observations indicate that ClinVar is at present an imprecise tool for assigning loss of function or pathogenicity to INSR variants, and illustrates the value of MAVEs in helping to refine classification.

**Figure 5:**
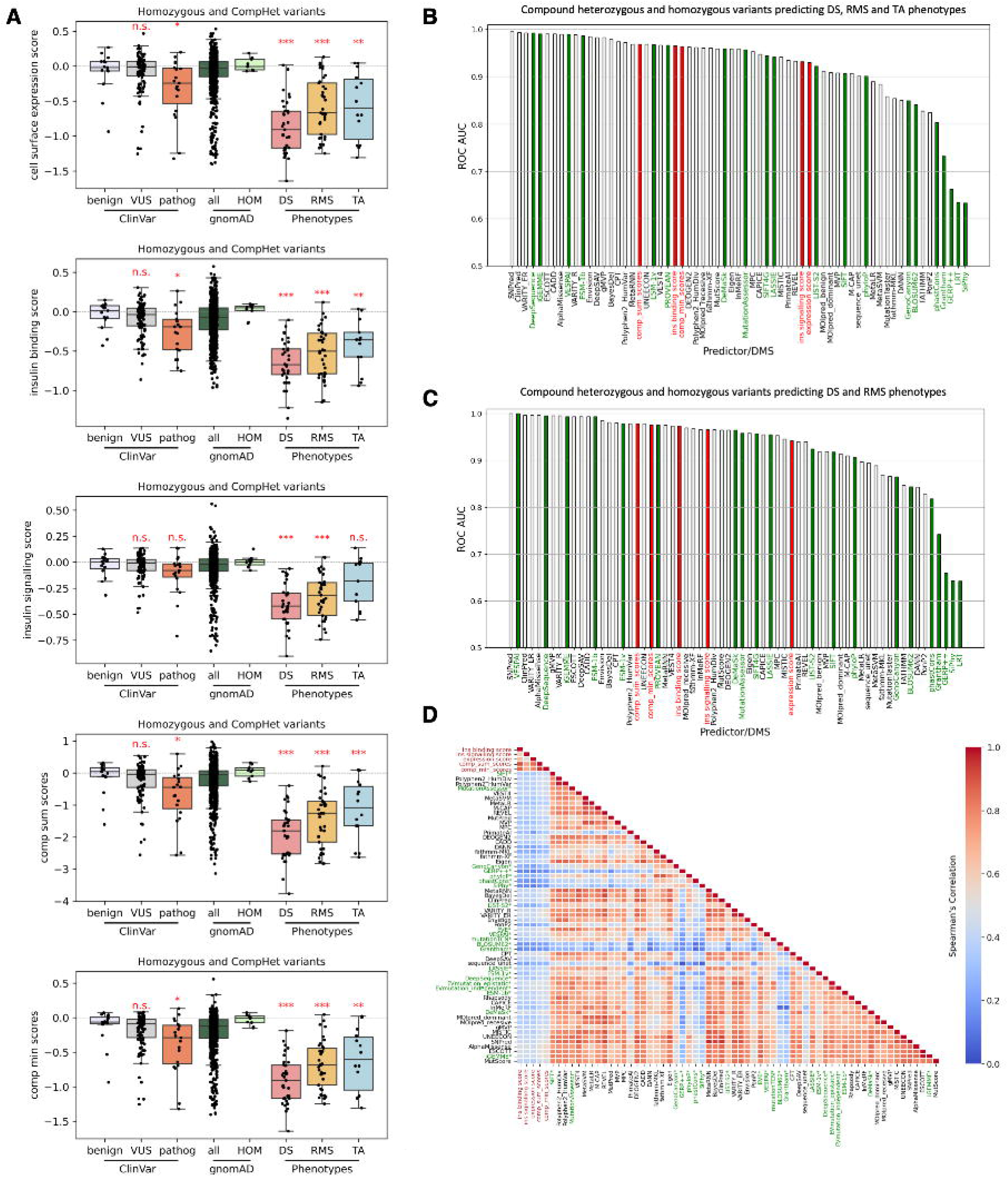
Evaluation of multiplexed assay results as an aid to genetic diagnosis. **A**, Boxplots showing expression, insulin binding, and signaling score distribution, as well as composite minimum or composite sum of scores for different groups of variants: ClinVar variants with any designation as benign or likely benign, as variants of uncertain significance (VUS), or as pathogenic or likely pathogenic are shown. Data for all variants and homozygous variants from gnomAD are also shown, as well as scores for curated pathogenic variants associated with Donohue syndrome (DS), Rabson- Mendenhall syndrome (RMS), or type A insulin resistance (TA). The plots contain scores for individual variants from gnomAD, as well as DS, RMS, and TA variants. See STAR Methods for average score computation for compound heterozygous variants. Mann Whitney U test was utilized to test if DS, RMS, and TA variants differ from gnomAD homozygous variants and if ClinVar VUS and pathogenic variants differ from ClinVar Benign variants. *, ** and *** denote p-values less than 0.05, 0.01 and 0.005 respectively. **B**,**C**, Receiver Operating Characteristic (ROC) analysis showing the Area Under the Curve (AUC) for all curated individual pathogenic variants for (**B**) DS, RMS, and TA and (**C**) only DS and RMS. The analysis compares predictions made using a set of supervised (light grey) and unsupervised (green) methods for variant effect prediction with those based on expression, insulin binding, and signaling scores alone, as well as composite minimum or sum of scores calculated in this study (red). Homozygous gnomAD variants were used as the set of benign variants. **D**, Spearman’s correlation matrix for scores from this study (red axis labels) and those from supervised predictors (black axis labels) and unsupervised predictors (green axis labels).

Diagnostic interpretation of INSR ectodomain variants is complicated by recessive inheritance, with clinical outcome depending on the composite effect of both INSR alleles. In keeping with this, 36 curated pathogenic variants and 122, 276, and 170 variants with severely reduced expression, insulin binding and signaling scores respectively were found in heterozygous form in gnomAD (**Figures 5A**; **S8C-E**). 68 variants in gnomAD showed severely reduced scores in all three assays (**Figure S8F**), and some loss-of-function variants were found at frequencies above 1/10,000 (**Figures S8C-E**). These may be valuable for future reverse genetic studies of INSR function.

To address the challenge of recessive inheritance of INSR-related severe IR in patient-level analysis, we calculated average function scores for pairs of homozygous or compound heterozygous variants associated with Donohue Syndrome (most severe), Rabson Mendenhall Syndrome (intermediate severity) and Type A insulin resistance (least severe). To enable this, we first adjudicated on clinical classification of all reported phenotypes to ensure consistency, including all pathogenic variants in ClinVar as well as all convincing pathogenic variants ascertained through two large genetic testing laboratories or previous clinical genetic reports (**Table S5**). We identified 88 unique extracellular missense variants as pathogenic in 99 different biallelic combinations, 43 of which were homozygous. Some extracellular missense mutations co-occurred with nonsense, frameshift or whole exon deletions in the extracellular domain, or intracellular domain mutations, to which we assigned low scores to reflect their anticipated effects (see Materials and Methods).

We found a clear correlation between average biallelic scores from the multiplexed assays and clinical syndrome, with the lowest scores in DS, intermediate in RMS and highest in type A insulin resistance (**Figure 5A**). Scores in all three groups were lower than for the group of all missense variants in the gnomAD repository, except for signaling by variants described in Type A insulin resistance. Scores in disease groups were strikingly lower than for the small number of presumed benign homozygous variants in gnomAD, but overlapped with scores for all variants in gnomAD, as expected for a recessive condition.

We then evaluated the ability of biallelic scores to predict severe insulin resistance, taking pathogenic mutations from the curated and adjudicated set identified in patients (**Table S5**), and defining homozygous variants in gnomAD as benign. A sophisticated array of variant effect predictors (VEPs) has been developed to aid in silico prediction of variant pathogenicity, and these continue to evolve with incorporation of artificial intelligence-based approaches. We thus compared performance of the current MAVE against a basket of VEPs, using both results from individual MAVE experiments (expression, binding and signaling) and of aggregated scores (lowest score across all three assays and sum of all three scores). Using receiver operating characteristics (ROC) analysis, we found near-perfect predictive performance of the current MAVE and the top VEPs , all with areas under the curve (AUC) of greater than 0.96 both for all pathogenic variants, and those seen only in severe recessive disease (**Figure 5B,C**). Among MAVE scores, the sum of scores for each assay type, the lowest score in any of the assays, and the insulin binding score performed best, followed by expression and signaling scores. Interestingly, despite the high predictive performance of MAVEs and VEPs, even the top VEP-derived scores showed only moderate correlation with MAVE data (range of correlation coefficients for the best performing MAVE score (sum of scores) and VEP scores 0.55 (AlphaMissense) to 0.16 (GRE++)), whereas VEPs often correlated strongly with each other (correlation coefficients up to 0.98; **Figure 5D**). This illustrates that the two types of approach access complementary information and may support each other in pathogenicity prediction.

#### Responsiveness of mutated receptors to antibody partial agonists

Experimental evidence at scale complementing in silico VEPS will help address the escalating diagnostic problem of VUS. However the translational aim of this study extended beyond this. By undertaking multidimensional assays (of cell surface expression, insulin binding and insulin-induced signaling) by the library of receptor variants, we were able to determine which variants are expressed at the cell surface yet show severely impaired signaling. It is these variants that may be amenable to activation by atypical agonists such as the anti-INSR monoclonal antibodies studied. We tested this directly by assessing antibody-induced signaling. The experiment was designed as for insulin stimulation, but used 10 mM antibodies in lieu of insulin.

Both monoclonal antibodies tested were confirmed to elicit a pattern of signaling overlapping heavily with that of insulin (**Figure 6A-D**). Several key areas of difference were apparent on the variant effect map, however, including but not limited to the αCT region noted previously. We next computed the difference between antibody-induced and insulin-induced signaling scores. To focus on variants with attributes most suggestive of a beneficial response to antibody stimulation in vivo, we constrained analysis to variants showing severely impaired insulin responsiveness (**Figure 6E**). Differential signaling scores are represented as Manhattan plots for each antibody (**Figure 6F,G**), with areas visually identified on variant effect maps as showing selective activation by antibodies boxed for reference. Overall we identified 1193 and 1141 variants poorly responsive to insulin but responsive or less impaired with respect to mAb 83-7 and mAb 83-14 stimulation, respectively. These were not exclusively found in areas forming insulin binding sites 1 and 2, but relative preservation of antibody stimulation compared to insulin was particularly striking in these areas. 69% of all variants with relatively preserved antibody stimulation were responsive to both monoclonal antibodies tested, with differential responses for the remainder only particular accounted for by residues forming parts of mapped antibody epitopes (**Figure 6H**). D734A and S350L, variants which were the focus of previous proof-of-concept studies, were confirmed to signal on antibody exposure, but were far from the most responsive variants to antibody stimulation. The list of variants generated in this analysis encompasses good candidates for future translational studies of humanised versions of the antibodies studied here (**Table S6**).

**Figure 6:**
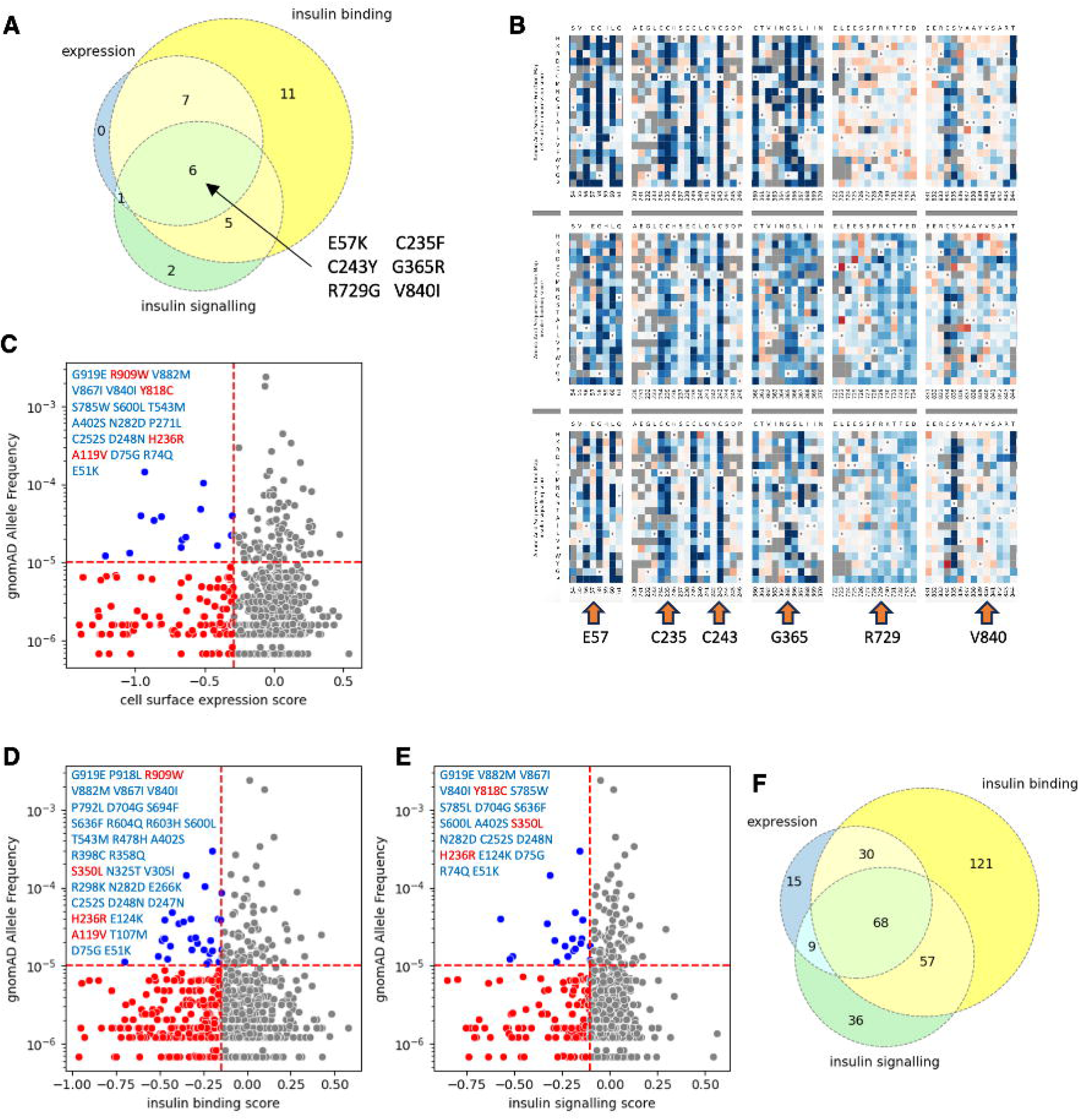
Functionally impaired variants in ClinVar and/or gnomAD datasets: **A**, Venn diagram comparing results for categorical loss of expression, insulin binding, and signaling determined by applying cut-offs from Figure 6A-C for variants classified uniquely as VUS in ClinVar. **B**, Heatmap excerpt of the 6 variants in the center of the Venn diagram in (**A**) showing loss of function in all three assays. **C,D,E,** Scatter plots showing allele frequencies for variants present in gnomAD v4.1.0(accessed 06/08/2024) against function scores for (**C**) cell surface expression, (**D**) insulin binding, and (**E**) maximal insulin signaling. The vertical dashed line indicates the cut-off scores derived as illustrated in **Figure S7A-C**. Variants with scores below these cut-offs are coloured red. The horizontal dashed line represents an arbitrary allele frequency of 10^-5, judged of potential utility for recall by genotype studies. Variants with scores below the function cut-offs but with allele frequencies above 10^-5 are coloured blue. These variants are listed on the plot. Variants identified and adjudicated as pathogenic in patients are coloured red **F**, Venn diagram for all variants listed in any of **C**, **D**, or **E**.

**Figure 7:**
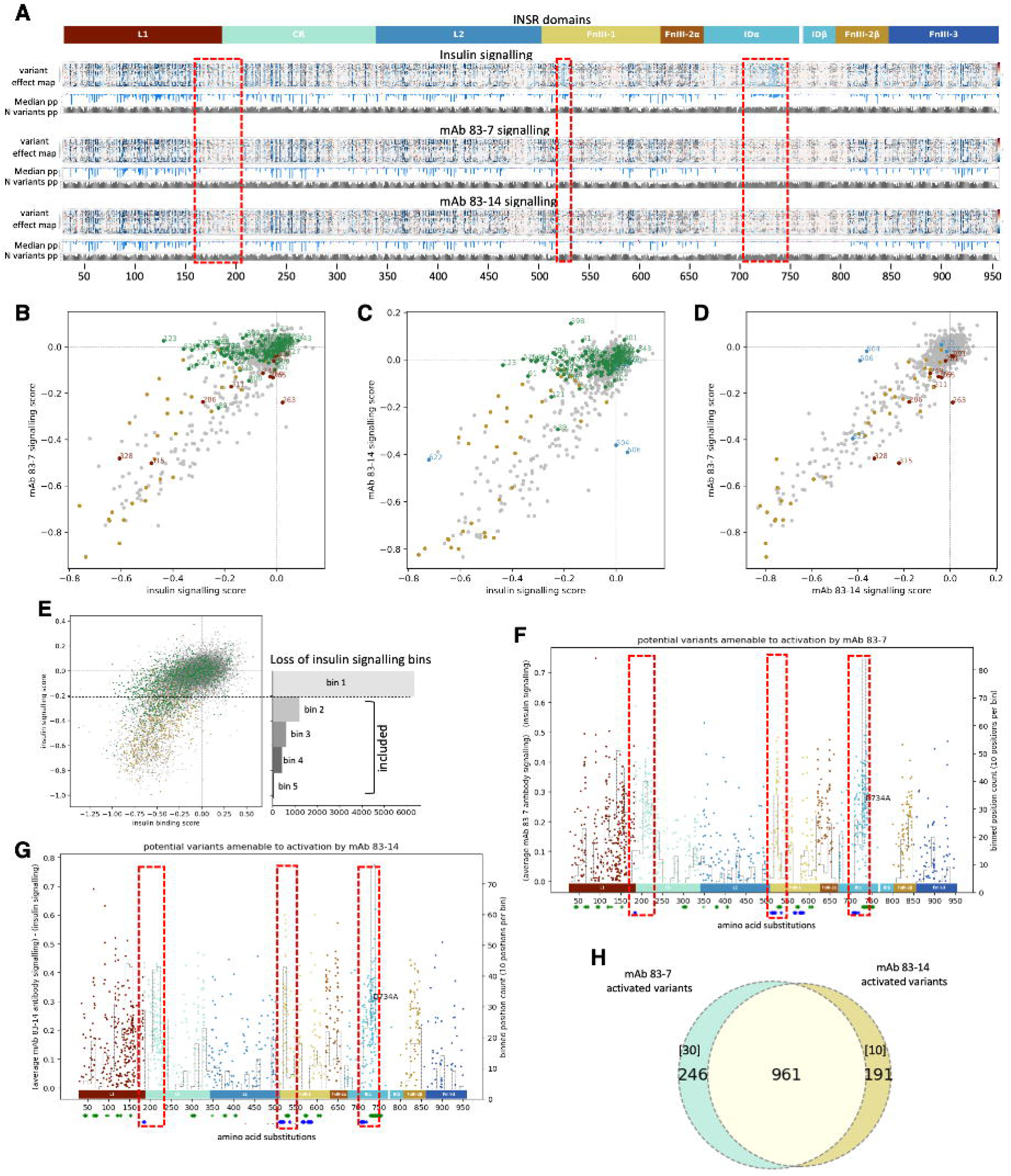
INSR variants with impaired insulin binding that are potentially activatable by monoclonal antibodies. **A**, Heatmap of variant scores for maximal signaling induced by insulin, mAb 83-7 or mAb 83-14, referenced to extracellular domain architecture. Variants that were not scored are coloured grey. ‘median pp’ denotes median variant scores per position and ‘N variants pp’ number of variants scored per position. Dashed red boxed indicate areas with visibly differential stimulation by insulin and antibodies. **B,C**, scatter plots of median signaling scores for (**B**) insulin and mAb 83-7 with epitope residues in dark brown or **(C)** insulin and mAb 83-14 signaling, with epitope residues in light blue. Cysteine residues involved in disulphide bond formation are coloured golden brown in both plots. **D,** Scatter plot of median scores for signaling in response to mAb 83-7 and mAb 83-14. **E**, Scatter plot of insulin binding and signaling variant scores. The horizontal box plot categorizes residues into 5 bins based on insulin signaling loss. Residues in the second bin or lower are classified as loss-of-signaling variants and used for Manhattan plots. **F**,**G**, Manhattan plots displaying differential signaling scores for (**E**) mAb 83-7 and (**F**) mAb 83-14 compared to insulin for all variants whose mutation impairs insulin signaling. Only variants with antibody signaling results from all three replicates are included in Manhattan plots, while only residues which both scored in all five insulin signaling replicates and which fall in bin 2 and below (illustrated in **E**) are plotted. Green or blue stars below X axes indicate site 1 and site 2 residues respectively. **H**, Venn diagram showing overlap between variants amenable to activation by mAb 83-7 and mAb 83-14, as plotted in Manhattan plots. The brackets show number of variants present in the respective epitope of each antibody.

## Discussion

The insulin receptor is of immense biomedical importance, playing a critical role in diabetes and its treatment, in the common disorders associated with insulin resistance, in growth, development,^3,4^ cancer, and beyond. Yet despite decades of intensive study , much remains to be learned about its complex insulin binding kinetics and signal transduction mechanism, and how its action goes awry in disease. Meanwhile for people with extreme insulin resistance due to biallelic mutations in the INSR gene, therapeutic options are poor.

Over 36 years since description of the first pathogenic INSR mutations^39,40^ around 70 pathogenic missense mutations have been described in the extracellular, insulin-binding portion of the receptor, and more continue to be found regularly. Functional studies, using various cell types and assays, have been published for only around 40 of these. We now describe a multidimensional MAVE of cell surface expression, insulin binding and peak insulin-induced signaling of around 14,000 such INSR variants, or 79% of all possible missense mutations. This offers a step change in functional understanding of insulin receptor variation of immediate diagnostic and translational value.

The current MAVE reproduces the large majority of findings from prior functional studies, especially for variants showing severe loss of function. Discrepancies were observed, however. In some cases they could be attributed to assay conditions (for example relating to “correction” of mutation effect by trypsin exposure), or to study of different receptor isoforms in the current MAVE (isoform B) and prior studies (not always explicitly documented). Some discordance may reflect uncertainty in MAVE results, but in many cases it is likely that previous assays are the major source of error. In nearly all cases these involved pairwise comparison of variant function against wild type receptor, often expressed from randomly integrated cDNA constructs, and linearity of readouts such as immunoblotting was commonly not demonstrated. In the current MAVE, in contrast, most variants were represented by more than one barcode/clonal cell line, in all cases with multiple biological replicates. Moreover all variants were compared simultaneously both against each other, and against thousands of wild type receptors, with a quantitative assay readout.

Comparing predictions based on the current study scores against a raft of in silico variant effect predictors (VEPs), we find that our data, like the best VEPs, performs outstandingly in predicting pathogenicity. Nevertheless ROC analysis relies on binary partitioning of variants into pathogenic and benign categories, and, as noted below, not all receptor attributes have been assayed in this study. By incorporating findings from further assays, it is likely that the ability of INSR MAVEs, and by extension other receptor tyrosine kinase MAVEs, to discriminate pathogenic mutations may be enhanced further. On the other hand, VEP performance is also improving. The current MAVE serves to validate performance of the best current VEPs in pathogenicity prediction, however, interestingly, correlation among scores from in silico predictors and empirical data is modest. It thus appears that MAVEs yield information complementary to in silico prediction alone.

Importantly, the current MAVE has value beyond tackling the VUS “roadblock” in diagnosis. First, we demonstrate the utility of MAVEs for functional mapping of antibody epitopes. More importantly, we exploit the multidimensional nature of our functional studies to inform future translational studies of insulin receptoropathy. Specifically, we aimed to stratify INSR variants for studies of novel receptor agonists by identifying which receptor variants are expressed at the cell surface and thus amenable to activation by non canonical ligands. By comparing antibody and insulin-induced signaling we further aimed to stratify mutations prospectively for future trials of humanised antibodies or other novel INSR agonists, including peptides and aptamers. Our dataset will aid early identification of infants with recessive INSR defects who might respond to INSR ligands with a distinct mode of receptor engagement to insulin. Very poor prognosis and short lifespan commonly seen in DS greatly constrains time available to assess new variants. We thereby tackle a key rate determining step in translational development of precision therapy for recessive receptoropathy.

The insulin binding and signaling assays we used, using either no insulin, or a single high insulin concentration, were not designed to detect variants conferring gain of insulin binding or signaling in the physiological insulin concentration range. This study did, however, identify several variants that increase maximal insulin binding and/or signaling, particularly clustered in regions known to contribute directly to insulin binding or to be involved in dynamic remodelling of the receptor on ligand binding. Further study of these and other possible gain of function variants using more physiological insulin concentrations in future will be of great interest.

Despite its large scale, our study has some limitations. First, around 20% of the full repertoire of missense mutations have not been studied, although in many cases inferences can be drawn from other substitutions of the same residue. A second limitation is the use of only a single very high insulin concentration. This reports only maximal response to receptor activation, or efficacy, and will miss changes in potency, assessed by clinically significant rightward shifts in the sigmoidal dose response curve to insulin. Moreover some variants that are convincingly pathogenic on genetic and clinical grounds show largely normal behaviours in this study and many prior studies even where dose response studies are undertaken. These include I146M^41^ and K487E^42^. This reflects the inability of simple assays to capture the full range of receptor attributes important for in vivo function, including dynamics of receptor recycling after ligand stimulation. For I146M and K487E, defects in receptor function were only unmasked on more complex cellular studies of receptor recycling kinetics^41,42^. It will be of great interest to use the cellular INSR library generated in this study to extend analysis to dose response relationships and recycling kinetics. Future work could further screen different ligands, different downstream pathways, or INSR expressed as a hybrid with the homologous IGF1R. Mixing of ligands with different fluorophore labels may also allow interrogation of binding co-operativity.

A final limitation arises from the cellular model used, combining Igf1r knockout and conditional endogenous Insr knockdown. This showed some evidence of low level residual endogenous Insr expression, which is likely to have narrowed the dynamic range of signaling and insulin binding assays. This will not have affected antibody stimulation and binding, as the antibodies used are human INSR-specific. For these technical and biological reasons the current signaling readout likely has greater specificity than sensitivity for detection of impaired signaling. For example, it will likely fail to identify variants for which the dose response curve is right shifted, but for which peak signaling at very high insulin concentrations (Emax) is normal. In use of our findings to aid genetic diagnosis this problem is mitigated by use of composite scores incorporating data from all three assay types. Nevertheless our findings are likely to be of greatest value for providing evidence for loss of function.

In summary, we report a large multidimensional MAVE of the extracellular INSR. This has high translational utility in monogenic insulin resistance, is an important starting point for more sophisticated studies of variant receptor attributes, informs on complex structure-function relationships of the INSR, and serves as a paradigm for study of other RTKs of high biomedical importance.

## Acknowledgements

RKS and GK are supported by the Wellcome Trust [grants 210752 and 207507]. RKS is additionally supported by the BHF Centre for Research Excellence Award III [RE/18/5/34216], and GK by UK Medical Research Council (MRC) University Unit programme MC_UU_00035/8. GVB & RKS are supported by Diabetes UK [grant 22/0006407]. CV has been funded via the PRISIS rare disease reference center, funded by the French Ministry of Health and Assistance-Publique Hôpitaux de Paris, and is a member of the European Reference Network on Rare Endocrine Conditions (EndoERN Project ID n° 739527). We are grateful to Ben Livesey and Joe Marsh (University of Edinburgh/MRC Human Genetics Unit) for advice on ROC analysis methodology, and thank Elisabeth Freyer and Michael Rennie (Flow cytometry facility, University of Edinburgh Institute of Genetics and Cancer) for support with extensive cell sorting. We thank Stephen O’Rahilly (University of Cambridge/MRC Metabolic Diseases Unit), Olivier Lascols (Department of Molecular Biology and Genetics, Saint-Antoine Hospital, Assistance Publique-Hôpitaux de Paris, Paris, France) and Drs Philip Gorden and Rebecca Brown (National Institute for Diabetes, Digestive and Kidney Disease, Bethesda, Maryland, USA) for access to INSR variant details and associated clinical data from their respective centres, and all clinicians referring patients for genetic studies.

## Author Contributions

Conceptualization: VA,RKS,GK; Data curation: VA, RK, HÇ, GVB, RK,CV; Formal analysis: VA, GVB; Funding acquisition: RKS,GK; Investigation: VA; Methodology: VA,GVB; Project administration: VA,RKS; Resources: VA,RKS,GK,GVB,KM; Supervision: RKS,GK; Validation: VA,GVB; Visualization: VA,GVB; Writing – original draft: VA,RKS,GK,GVB; Writing – review and editing: all co-authors

## Supplemental information

Supplemental Figures S1 (Related to Figure 1); S2 (Related to Figure 1); S3 (Related to Figure 2); S4 (Related to Figure 2); S5 (Related to Figure 3); S6 (Related to Figure 4); S7 (Related to Figure4)

Supplemental Tables S1 (Related to Figure 1); S2 (Related to Figures 2-7); S3 (Related to Figures 2-7); S4 (Related to Figures 2,4); S5 (Related to Figure 5); S6 (Related to Figure 7); S7 (Related to STAR Methods) (all but one too large to fit into pdf)

Supplemental Reference List (To be completed: All PMID listed in Supplementary tables for first submission)

## STAR* Methods

### EXPERIMENTAL MODEL AND STUDY PARTICIPANT DETAILS

#### Curation of prior clinical, genetic and functional data

Pathogenic INSR mutations were ascertained from ClinVar Miner^43^ (accessed 2.8.2024), from curation of published reports, and from unpublished diagnostic testing results from national referral centres for severe insulin resistance in Cambridge, UK and Paris, France (**Table S4)** . All available clinical data were examined and subjective adjudication was made on clinical diagnosis, irrespective of reported diagnostic labels, based on reported biochemical severity of insulin resistance, birthweight and other phenotypic features, and longevity. Donohue syndrome and Rabson Mendenhall syndrome cannot be demarcated precisely, with longevity (1-2 years for Donohue syndrome, >10 years for Rabson Mendenhall Syndrome) a key discriminator. Prior published functional studies of INSR variants, both naturally occurring and generated for experimental purposes, were curated and assessed. A wide variety of functional studies was reported, complicating simple comparison of findings, whose details are summarised in **Table S4** . For each of expression, insulin binding and signaling, findings were stratified into severe loss of function (approximately <10% wild-type function), moderate loss of function (>10% wild-type function) and normal function. INSR variants in gnomAD were drawn from gnomAD v4.1.0, accessed 6.8.2024).

#### Generation of R-MmINSR KD cells containing single landing pads

Murine embryonic fibroblasts (MEFs) derived from the Igf1R^-/-^ mouse line^44^ were infected at low multiplicity of infection with lentivirus encoding concatenated shRNAs targeting the Insr gene^14^, and exposed to 500ug/ml hygromycin-B for 2 weeks. Clonal cell lines were isolated by limiting serial dilution and screened for GFP expression upon DOX addition. Four clones demonstrated strong inducible knockdown of endogenous Insr, one of which was used for all subsequent experiments.

A previously described Tet-inducible BxB1 DNA recombinase landing pad (Tet-coBxb1-2A-BFP_IRES- iCasp9-2A-Blast_rtTA3, Addgene 171588) was introduced into R-MmINSR KD cells using the Lenti-X Packaging Single Shot (VSV-G) (Cat 631275) system. Briefly, 7μg of pLenti_Tet-coBxb1-2A-BFP_IRES- iCasp9-2A-Blast_rtTA3 lentiviral vector in 600ul water was mixed with Lenti-X Packaging plasmid and incubated at room temperature for 10 minutes. The mixture was transferred dropwise onto cultured Lenti-X 293T cells on a 10 cm plate and incubated at 37 °C in 5% CO2. Medium was changed the next day and supernatant collected after 48 and 72 hrs and pooled. Collected medium was centrifuged (300g, 5 min), and the supernatant filtered through a 0.45μm filter to remove debris. The landing pad construct expressed doxycycline-inducible blue fluorescent protein (BFP) and a Blasticidin resistance gene, which were used to confirm landing pad insertion. 100μl to 1ml lentiviral supernatant was used, and assessment of BFP expression 48 hours after infection was used to identify lines with an MOI <1. After treatment with 6 μg/ml Blasticidin for one week, cells with the highest BFP fluorescence were sorted into single cells in 96-well plates using a BD FACS Aria II (405- 450/50 nm laser). Surviving clones were transferred into a 24-well plate and later 6-well plates. 400,000 of the 3 selected clonal lines were seeded on a 6 well plate and transfected with 2.5ug of attB-miRFP670 using Lipofectamine 3000. The medium was changed and 2ug/ml Puromycin added 48 hours after transfection. Cells were assessed for recombination after a week of antibiotic selection with a BD LSR Fortessa flow cytometer. One clone showing uniform loss of BFP and gain of miRFP670, indicative of a single landing pad was selected for use in subsequent experiments.

### METHOD DETAILS

#### General reagents

Enzymes were purchased from New England Biolabs unless otherwise stated. Oligonucleotides were from IDT and are listed in **Table S7**. Igf1r^-/-^ mouse embryo fibroblasts (MEFs) were obtained from the Cosgrove laboratory CSIRO, Adelaide, Australia^44^. MEFs were cultured in Dulbecco’s modified Eagle’s medium (Thermofisher, 41965039) supplemented with 10% fetal bovine serum and Penicillin-Streptomycin (50 I.U./ml). Cells were passaged by trypsinization using 0.25% trypsin-EDTA (Millipore-Sigma) and tested negative for Mycoplasma before use.

#### Plasmid construction

The open reading frame of the B isoform (exon 11+) of the human INSR gene (UniProt P06213-2) was subcloned from pCDNA5/FRT/TO/hINSR^14^. attB_INSR_P2A_PuroR plasmids were created by restriction cloning, combining amplicons from attB_sGFP-PTEN-IRES-mCherry-P2A-PuroR (a gift from Douglas Fowler, University of Washington) amplified with primers VA01//VA02, and pCDNA5/FRT/TO/hINSR amplified with primers VA03//VA04 and VA05//VA06. To permit barcode cloning into the attB_INSR_P2A_PuroR plasmid, new restriction sites were created by digesting the plasmid with AfeI, dephosphorylating with Shrimp Alkaline Phosphatase (rSAP, M0371), and ligating an amplicon containing SbfI and MluI sites. The amplicon was made by annealing 10mM of phosphorylated VA07 and VA08 primers before purification, ligation into the plasmid with T4 DNA ligase, and transformation into XL-10 gold cells. Correct insert orientation was confirmed by Sanger sequencing.

#### Plasmid barcoding

VA9 30mer barcoding oligos flanked by Sbf1 and AscI sites were synthesized and PAGE-purified before VA10 primer annealing and extension using Phusion HiFi PCR master mix (M0531S). 10 separate 50ul reactions, each using 0.5uM primer and 0.1uM barcoding oligo were mixed, purified using two MinElute PCR Purification columns (QIAGEN, 28004) and quantified with Nanodrop. 32ng double stranded barcoding amplicon was digested overnight at 37℃ in a 500ul reaction containing10ul SbfI-HF and 10ul AscI, and purified with two MinElute columns. In a separate reaction, 7ug attB_INSR_P2A_PuroR plasmid was digested for 2 hours in a 300ul reaction supplemented with 5ul Sbf1-HF and 5ul MluI-HF followed by 30 minutes dephosphorylation with Shrimp Alkaline Phosphatase (rSAP, M0371) and purification with QIAquick PCR Purification columns (QIAGEN, 28104). Digested barcoding amplicon was cloned into digested attB_INSR_P2A_PuroR plasmid with T4 DNA ligase in 5 separate 3ml reactions (each containing 600ng digested plasmid, 5.25ng digested barcoding amplicon, 300ul T4 DNA ligase buffer and 5ul T4 DNA ligase) with 4 hours room temperature incubation. Reactions were supplemented with 2.5ul each of the isoschizomers MluI-HF and AscI to remove any background from plasmid self-ligation. Ligation of MluI-HF and AscI- derived fragments abolishes the recognition sequences. 15ml Qiagen PB buffer was added to each reaction, then eluted in 15ul using a DNA Clean & Concentrator column (Zymo Research, D4013). Eluted samples were pooled and digested by MluI-HF and AscI for 1 hour to remove residual background. 9ul of this reaction was transformed into 100 of XL10-Gold Ultracompetent Cells in 10 separate reactions (Agilent Technologies, 200315). Before plating, transformation mixes were pooled in a total volume of 10ml, 11ml sterile water was added and 400ul of the mixture was quickly spread on 52 150mm agar plates with 100ug/ml ampicillin, using glass beads. Dilution plates were made with 10ul, 25ul and 50ul of the mixture to allow estimation of transformation efficiency and number of barcoded plasmids. All plates were incubated at 37℃ overnight. Colonies were scraped, pooled, and aliquots stored as pellets at -80℃.

#### Multiplexed codon mutagenesis

The library generation and sequencing workflow is outlined in **Figure 1**. First, previously described primer design software^45^ was used to generate a pool of 60mer oligonucleotides, consisting of 928 subpools, each including an NNS codon at one of the 928 codons of the extracellular receptor, flanked by 28 and 29bp of template-homologous sequence before and after the mismatched codon, respectively. The NNS codon encodes all 20 amino acids and one stop codon. To minimise unevenness of mutagenesis, oligonucleotides were organized into 47 pools, each covering 20 sequential codons, and separate reactions were run for each. Each oligonucleotide mixture was phosphorylated by adding 20ul 10uM oligonucleotides to 2.4μl T4 Polynucleotide Kinase Buffer, 1ul 10 mM ATP, 1ul T4 Polynucleotide Kinase (10 U/μL) into a PCR tube and incubation at 37°C for 60 minutes. Phosphorylated oligos were stored at -20°C and diluted 1:300 in nuclease-free H_2_O on the day of mutagenesis. 7μl 100μM universal secondary primer (VA11) was phosphorylated similarly in a final reaction volume of 30μl, and diluted 1:20 on the day of mutagenesis.

We applied a reported nicking mutagenesis strategy^19^ to construct the INSR variant library. The barcoded attB_INSR_P2A_PuroR plasmid library was freshly purified from a bacterial pellet using the Qiagen Mini-Prep Kit (Cat. 27106X4) for mutagenesis before column purification, elution in 10µL nuclease-free H_2_O and transformation into 100ul XL10-gold ultracompetent cells (Agilent, 200315). Transformation mix volumes were adjusted to 850μL with sterile LB medium and 800μL spread on two 150mm agar plates containing 100mg/ml Ampicillin. Serial dilutions from the remaining 50ul reaction mixture were prepared to permit calculation of transformation efficiencies. After overnight incubation at 37°C, colonies were scraped into 10mL LB at an estimated bottleneck of 130,000 (whole library) or 2750 (individual mutagenesis blocks) colony forming units to limit library size. 3ml scraped cell suspensions were prepared using a Qiagen Mini-Prep Kit and 2ug purified plasmid DNA from each block was pooled to prepare the final barcoded, mutagenised INSR library.

#### PacBio sequencing of INSR variant library for barcode-variant mapping

PacBio sequencing was used to phase barcodes and mutations through long sequence reads spanning barcode and INSR extracellular domain sequence. To eliminate strand exchange during PCR, PacBio sequencing templates were prepared from purified plasmid library by SbfI-HF and PspOMI digest. Digested product containing 3kb template fragment (barcode and ectodomain sequence) and 5kb plasmid backbone was cleaned using the QIAquick PCR Purification Kit and submitted to University of California, Davis PacBio Sequencing Services for size selection with BluePippin (Sage Science) followed by library prep and sequencing on a single SMRT cell with Sequel II, using 30-hour movie collection times. The Sequel II system performs on-instrument data processing and delivers accurate long reads (HiFi reads) through building circular consensus sequences (CCSs) from subreads (raw sequencing files available in GEO; accession number pending).

We processed HiFi reads using alignparse^46,47^ version 0.5.0, which applies minimap2^48^ version 2.24 for long read alignment to identify and call mutations in the INSR sequence and to phase them with barcodes. 3,568,566 of 4,016,907 HiFi reads (89%) mapped to the target (available in GEO; accession number pending), and the output was used to generate a codon-variant lookup table. We first retained mapped HiFi reads with sequencing accuracy reported by the PacBio ccs program in both INSR ectodomain sequence and barcode to be at least 99.99%. 70% of all reads passed these filters. As PacBio sequencing produced on average 19 reads per amplicon (**Figure S1B**), we next calculated the empirical accuracy of retained reads^47^ to assess the reliability of barcode-variant phasing, i.e. how often the reads linked with a specific barcode are identical. We observed 99.4% empirical accuracy for each HiFi read, excluding reads with indels. Since 96% of the barcodes were associated with two or more HiFi reads we generated a cumulative consensus of HiFi reads within these barcodes, so that consensus accuracy would exceed calculated empirical accuracy for individual reads. Barcodes differing by only two bases but associated with different variants were omitted from the codon-variant table. Selection criteria were designed to exclude barcodes with lower CCS read counts, while retaining those with higher read counts. As a result, 52 conflicting barcodes, mostly with single reads, were removed. Barcodes associated with insertions, deletions or more than one variant were also excluded. Overall 126,272 barcodes were retained, of which 80,972 associated with 15,996 missense variants, 3,539 barcodes with 813 nonsense variants, 4,023 and 41,232 barcodes with synonymous variants and wild type sequence, respectively (**Figure S1B-D; Tables S1,2**). The code used to generate the final barcode-variant table is available on https://data.mendeley.com/preview/4sm82c4bb5?a=9097473b-604f-48c1-b0b5-8d97042c0775 and includes additional plots illustrating library structure.

#### Transfecting cells with the INSR mutation library

Landing pad-containing R-MmINSR KD cells at 80% confluence were trypsinised and pelleted at 300g for 4 minutes at room temperature, washed with 20mL PBS, counted, and resuspended in OptiMEM at 1.17x10^8^ cells/mL. In a separate tube, 3ml of cells (3.5x10^8^ cells) were mixed with 2.8mg plasmid library resuspended in 500ul nuclease-free H_2_O. The mix was loaded on a MaxCyte CL-1.1 processing assembly (PA) using a 10ml luer-lock syringe, and electroporated twice using the MaxCyte GTx 0-6 setting. The electroporated cells were immediately unloaded using a syringe and the PA washed twice with 3.5ml OptiMEM to collect remaining cells. Cells were allowed to recover at 37°C for 40 minutes before 75mL warm growth media was added. The cell suspension was introduced to 15 5- layer T175 flasks to a density of 4.7x10^8^ cells/175cm^2^ and incubated under standard conditions. Selection with 1.5ug/ml puromycin was started 48 hours after transfection and the medium was changed every 2-3 days until colonies appeared and grew to high confluency for harvesting. Harvested cells from all flasks were pooled and aliquoted for cryopreservation.

#### Preparing fluorescent insulin and antibody conjugates

We used the Alexa Fluor 647 NHS Ester (Thermofisher, A37573) kit to label recombinant human insulin (Sigma, 91077C). Immediately before use, the amine-reactive compound was dissolved in anhydrous dimethylsulfoxide (DMSO) to 10 mg/mL and insulin in 0.1 M sodium bicarbonate buffer [pH 8.3] to prepare 10mg/ml insulin solution. 100ul reactive dye solution was slowly added to 266ul of this insulin solution and the reaction incubated for 1 hour at room temperature with continuous stirring. Labelled insulin conjugate was recovered by dialysis using Slide-A-Lyzer MINI Dialysis Devices, 3.5K MWCO (Thermofisher, 69550). 83-7 and 83-14 antibodies (gifts from Prof. Kenneth Siddle) were labelled with AF647 dye using Zip Alexa Fluor Rapid Antibody Labelling kit (Thermofisher, Z11235) following the manufacturer’s protocol.

#### Assaying insulin or antibody binding

Cells were recovered from liquid nitrogen, expanded for at least two passages, washed twice with PBS and serum starved (DMEM without FCS) for 16h. 1.5x10^8^ cells were used for each replicate. Cells were washed once with PBS, detached with trypsin, quenched with an equal volume of growth medium, and centrifuged at 400g for 4 minutes before washing with FACS buffer and resuspension at 1x10^7^/ml in FACS buffer. Cells were incubated with AF647_insulin (0.216 mg/ml), AF647_83-7 or AF647_83-14 (1mg/ml) conjugates for 1h on ice with periodic mixing. Stained cells were spun, washed 4 times with FACS buffer and fixed in 3% methanol-free PFA (43368, ThemoFisher) for 10 minutes on ice. Finally, cells were spun down, washed once with PBS, and resuspended in PBS before sorting on a FACS Aria II within 48 hours.

#### Assaying insulin- or antibody-induced Akt phosphorylation

Cells were prepared as for binding assays until quenching of tryspsinised cells with an equal volume of warm GM, this time containing 100nM human recombinant insulin (Sigma, 91077C) or 10nM anti-INSR antibody. After 10 minutes incubation at 37°C, cells were centrifuged at 400g for 4 minutes and washed with FACS buffer before fixing with 4% methanol-free PFA (Thermofisher, 28908) for 10 minutes on ice with periodic mixing. Fixed cells were washed once with FACS buffer and resuspended in fc block (FACS buffer with 10% FCS) containing 0.1% saponin and incubated on ice for 15 minutes to permeabilise cells and block Fc receptors. Fixed cells were spun, resuspended at 10^7^ cells per ml in FACS buffer containing 0.1% saponin, and stained with AlexaFluor 647-conjugated anti-Phospho-Akt (Ser473/474) antibody (Cell Signaling, 4075) at 1/200 dilution for 1h on ice with periodic mixing. Stained cells were spun, washed 4x with FACS buffer and fixed with 3% methanol-free PFA for 10 minutes on ice. Finally, cells were spun, washed once with PBS, and resuspended in PBS before sorting on a FACS Aria II within 48 hours.

#### Validation of FACS-based assays of INSR function

To validate FACS-based assays of INSR expression and function, we used non transfected cells, and cells transfected with either wild-type (WT) INSR or the mutant INSR library. Both monoclonal antibodies used robustly detected surface INSR expression in WT and library cells compared to untransfected cells (**Figures 1C**, **S1F**). Similarly, binding of labelled insulin produced comparable profiles for WT and library cells which were clearly distinct from the profile of untransfected cells, although some insulin binding was detected in control cells, consistent with low level residual expression of murine Insr.

A smaller dynamic range was observed in assays for phosphorylated AKT (pAKT) (**Figures 1C**, **S1F**), as expected for a labile intracellular antigen. Shifts in pAKT FACS profile were seen on stimulation by insulin, 83-7 or 83-14 antibodies for both WT and library cells, but some pAKT was also detected on insulin stimulation of untransfected cells. The lack of pAkt detection in untransfected cells stimulated with human INSR-specific antibodies further supports this being due to binding to endogenous mouse Insr, in keeping with the residual insulin binding noted above. We also observed that overexpression of either WT or mutated human INSR induced some AKT phosphorylation even without stimulation. However, despite these limitations of the cellular system used, we consistently observed much stronger binding of and signaling by insulin and antibodies than background.

#### Variant library FACS sorting and barcode detection

A BD FACS Aria II sorter was gated for singlets, and cells were sorted into four bins by fluorescence in the 640-670/14A channel. At least 13 million cells were collected per bin. Sorted cells were collected by centrifugation and stored at -20°C before gDNA extraction and library preparation.

Genomic DNA was prepared from cells in each bin by phenol-chloroform extraction. Cell pellets were resuspended in 500μl lysis buffer (100mM Tris [pH 8.5], 0.5mM EDTA, 0.2% SDS and 200mM NaCl, 25ug RNAase A) and incubated at 37°C for 1 hour before adding 700ug proteinase K (New England Biolabs, P8107S) and incubated at 56°C overnight. An equal volume of Phenol:Chloroform:Isoamyl (PCI) alcohol 25:24:1 was added to each reaction, vortexed vigorously and centrifuged at high speed for 5 minutes. The aqueous layer was transferred to a clean 1.5ml tube, mixed with equal amount of PCI, vortexed vigorously and centrifuged at high speed for 5 minutes. The aqueous layer was again transferred into a new tube and DNA precipitated with ethanol and resuspended in 400ul nuclease-free H_2_O. Because extracted gDNA was viscous, samples were sonicated on a Diagenode Bioruptor Plus on high power with 30 seconds on/off for 5 cycles.

PCR was performed to amplify gDNA barcodes. To avoid amplifying barcodes from any persisting extragenomic INSR plasmid, the reverse primer was designed to anneal to a gDNA sequence downstream from the plasmid integration site. For every bin, we distributed purified gDNA across multiple 25μL PCR reactions. Each included 12.5μL Phusion high-fidelity PCR master mix with HF buffer (NEB, M0531S), 1μg genomic gDNA, 10μM reverse primer with Illumina P5 sequence (VA12), and 10μM indexed forward primer including an Illumina P7 sequence (VA13) binding upstream from the barcode. PCR began with a step of 96°C for 2 minutes and ended with 72°C for 1 minutes, with 24 amplifcation cycles (96°C for 30 seconds; 65°C for 30 seconds; 72°C for 40 seconds) in between.

PCR products for each bin were pooled and 300ul purified with MinElute PCR Purification Kit and resuspended in 50ul of water. The eluate was size selected (expected size of 283nts) on a 2% agarose gel (Monarch DNA Gel Extraction Kit, T1020S) and quantified by Bioanalyser DNA1000 chip. Quantified samples were mixed together in equimolar ratios and submitted for 50bp single end sequencing using a custom sequencing primer (VA14) on an Illumina NextSeq. Demultiplexed reads were trimmed using trimmomatic software such that only 30nt barcode sequences was left. Sequences then aligned to the barcode sequence library determined from PacBio sequencing by Enrich2, yielding a count of the number of times each library barcode was sequenced in each FACS bin. Raw illumina sequencing and processed files are available on GEO. Scripts are available at https://data.mendeley.com/preview/4sm82c4bb5?a=9097473b-604f-48c1-b0b5-8d97042c0775

#### Generating G333Q, D627A and D743A single substitution cell lines

Primers (Key Resource Table) were designed to generate substitution mutations in attB_INSR_P2A_PuroR plasmid using the Q5^®^ Site-Directed mutagenesis kit following manufacturer’s recommendations. Products were amplified in E. coli strains and substitutions confirmed by whole plasmid sequencing. 10ug of variant plasmids were transfected into to 10*10^6 landing pad-containing R-MmINSR KD cells using MaxCyte PA R-50x3 before selection and study by flow cytometry.

### QUANTIFICATION AND STATISTICAL ANALYSIS

#### Calculation of variant function scores and false discovery rates

The puromycin selection marker is located downstream from the INSR gene, with a P2A self-cleaving sequence in between. Any stop codons in the INSR gene would stop translation of the Puromycin resistance gene and therefore cells carrying stop codons will be removed during antibiotic selection after transfection unless stop codon readthrough occurs. We expected no reads for such barcodes on Illumina sequencing but observed reads for 7% of the possible stop codon barcodes. We considered the possibility that these barcodes represent erroneous barcode-variant assignments in PacBio data analysis. To test this, we compared the counts of Illumina sequencing reads representing different classes of variants in FACS based assays. We found that barcodes associated with stop codons show greatly depleted read counts in the FACS data, compared to barcodes associated with wild-type, synonymous or missense variants. This strongly suggests that these barcodes represent bona fide stop codons that were only partially depleted during puromycin selection, perhaps thanks to stop codon readthrough, and supports the accuracy of our barcode-variant phasing approach.

We used Enrich2^49^ to count number of occurrences of each barcode in each bin. Enrich2 configuration files are available at https://data.mendeley.com/preview/4sm82c4bb5?a=9097473b-604f-48c1-b0b5-8d97042c0775. We then used a previously published method for calculating barcodes and variant scores with some modifications^50^. The count for each barcode in a bin was divided by the sum of counts recorded in that bin to obtain the frequency of each barcode (F_b_) within that bin. This calculation was repeated for every bin in each replicate experiment.

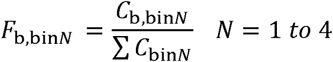

Barcodes with fewer than 150 reads across 4 bins in each experiment were filtered out. Next, for each replicate experiment, a weighted average (barcode score) was calculated for each barcode (W_b_) using the following equation:

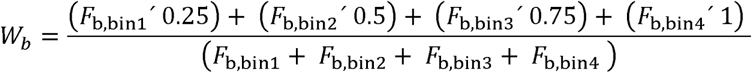

Finally, for each experiment, a score for each variant (*S_v_*) was obtained by dividing the mean of barcodes weighted average of each variant to mean of wild type barcodes weighted average as follows:

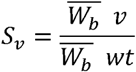

To quantify confidence that a variant’s score differs from the score of wild-type INSR, we used a bootstrapping approach to calculate, for each variant, a false discovery rate (FDR). This approach assigns FDR scores to variants associated with any number of barcodes, while avoiding assumptions about the distribution of scores and their measurement errors.

For variant *v* with *k* barcodes across all biological replicates, barcode score *B_V_* was calculated for each variant as:

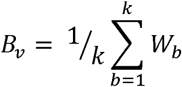

We then generated weighted averages of 10,000 randomly sampled sets of *k* wild-type barcodes *W_wt_* and calculated the wild-type barcode score *B_wt_* of each set as follows:

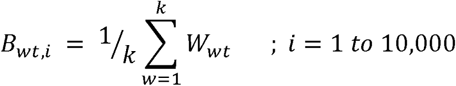

Finally we computed the false discovery rate *FDR_v_* for variant *v* from:

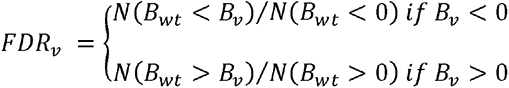

Where *N(B_wt_ < x)* is the number of times where *B_wt_ < x* across the set of 10,000 bootstrapped wild-type scores. A pseudocount of 1 is used if *N(B_wt_ < B_v_) or N(B_wt_ > B_v_)* is equal to 0.

#### Computation of biallelic function scores

Nonsense, frameshift or whole exon deletions, for which no empirical function scores were generated, were assumed to be complete or near complete loss of function variants, and were thus assigned the lowest high confidence score seen in the MAVE (for W516R); intracellular mutations, whether missense, frameshift or nonsense, were assigned this score multiplied by 1.5, representing a punitive factor to reflect the well documented dominant negativity of such mutations^51^.

## ADDITIONAL RESOURCES

Fully analysed data will also be interrogatable through a user-friendly web interface (www.xxxx)

## Supplemental Figure Legends

**Figure S1 (Related to Figure 1): Insulin Receptor Structure and Validation of Variant Library and Functional Assays. A,** Schematic of INSR with coloured domains (left) and cryoEM structure of the ectodomain bound to 4 insulin molecules (PDB 6PXV) coloured by domain using the same colour scheme. **B**, Distribution of PacBio circular consensus (CCS) reads per barcode, with median of 19 reads per barcode. **C**, Distribution of barcodes per variant. **D**, Mutations per variant showing that most INSR cDNAs in the plasmid library have either one amino acid change (a missense mutation), or no amino acid change. **E**, Frequencies of missense, synonymous and stop mutations in the plasmid library. **F**, FACS profiles for insulin or antibody binding (left panel) or stimulated signaling (right panel) for null (grey), wild type (blue) and library cells (red).

**Figure S2 (Related to Figure 1): Overview of massively parallel assay results and replication A**, INSR extracellular domain architecture with variant-function heatmaps for each replicate of insulin, mAb 83-7 and mAb 83-14 binding and signaling assays. **B**, Pearson’s correlation matrix showing coefficients of variation for all pairwise replicate combinations. Correlation coefficients (ρ) are annotated inside heatmap cells. **C**, Principal Component Analysis (PCA) of replicates.

**Figure S3: Extended data related to Figure 2. A**, Scatter plot of individual variant scores for insulin binding and cell surface expression. Site 1 residues are shown in green and cysteine residues involved in disulphide bond formation are coloured golden brown. **B,** Box plot comparing cell surface expression and insulin binding variant score distributions for non-cysteine residues with cysteine residues crucial for disulphide bond formation. **C**, Structure of 4 insulin-bound INSR ectodomain (PDB: 6PXV) with residues coloured by median cell surface expression score per position. All cysteine residues are shown in space filling style. **D**, Scatter plot of individual variant scores for mAb 83-7 binding, with dark-brown coloured outliers, and mAb 83-14 binding, with light-blue coloured outliers. Cysteine residues involved in disulphide bond formation are coloured golden brown for reference. **E**, Scatter plot of median scores for insulin binding and cell surface expression. Insulin binding site 2 residues are shown in blue and cysteine residues are coloured golden brown for reference. The inset boxplot shows expression and insulin binding scores for insulin binding site 2 residues. The Wilcoxon signed-rank test was utilized to determine if insulin binding and expression scores for site 2 residues differ from wildtype. Asterisks on the box plots indicate significance level. *, ** and *** denote p-values <0.05, <0.01 and <0.005 respectively. **F**, Scatter plot showing median insulin binding versus cell surface expression scores, with site 1 residues highlighted in green. The horizontal box plot stratifies residues into 5 bins based on expression loss. Residues in the second bin or lower are classified as loss-of-expression variants and were coloured white when colouring the structure in Figure 2G by insulin binding score.

**Figure S4 (Related to Figure 2): Comparison of massively parallel insulin binding scores with published alanine scanning studies of INSR ectodomain A**, Correlation of insulin binding scores from this study with ratios of mutant to wild-type insulin dissociation constants (Kd) from reported alanine scanning studies^29,30,32^, highlighting variants with FDR <0.05 in red. **B,** Correlation of median insulin binding scores per position from this study with these ratios. In **A** and **B** Kd ratios below the authors’ suggested confidence threshold for alanine scanning data are indicated by a shaded area. **C,** Heatmap excerpt displaying variant scores for the majority of residues with available alanine scanning data.

**Figure S5: Extended data related to Figure 3.** INSR residues identified to result in gain of function upon substitution mapped to the cryo-EM structures of the various insulin-bound states. Monomers are coloured different shades of grey. Insulin is represented by green ribbon structures. Yellow depicts residues where multiple substitutions increase insulin binding. Blue depicts residues where a unique substation increases binding. Structures are of the A isoform but residues highlighted are numbered according to the B isoform pro-receptor for consistency. Some residues highlighted may be involved in interactions with parts of the receptor not resolved in available structures. **A** and **B,** Two insulin bound (green ribbons) INSR (PDB: 7STJ (asymmetrical), and 7STH (symmetrical)). These structures are the A isoform of the mouse orthologue, but for consistency residues highlighted are numbered according to the human B isoform pro-receptor with the corresponding mouse residue in brackets. Highlighted sections are rotated counterclockwise 90°. **E)** Saturated, four insulin bound INSR (PBD: 6PXV). The highlighted section is rotated counterclockwise 90° to the full view.

**Figure S6 (Related to Figure 4): Residues whose mutation confers gain of binding or signaling. A**, Detail of scatter plot of median expression scores against insulin signaling scores, focused only on gain of signaling. **B**, Scatter plot of gain-of-insulin-signaling scores with FDR < 0.01 against expression scores for individual variants, for variants with scores in at least 3 replicates. **C–G,** INSR residues identified to result in gain of function upon substitution mapped to available cryo-EM structures of the various insulin-bound states. The two monomers are coloured different shades of grey. Insulin is represented by green ribbon structures. Yellow depicts residues where multiple substitutions result in gain of insulin binding. Blue depicts residues where a unique substation results in gain of binding. Orange depicts residues where a unique substitution confers gain of insulin-stimulated signaling. Purple depicts residues where substitution increases both insulin binding and signaling. Some of the flexible ID domain is not observed in the structures, and residues highlighted may be involved in interactions with parts of the receptor not resolved in available structures. **C,** Unbound, human INSR B-isoform (PDB: 8U4B). The highlighted section is rotated counterclockwise 45° to the full view. **D,** Single insulin bound (green ribbon) mouse INSR isoform A (PDB: 7STI). The highlighted section is rotated counterclockwise 90°. **E** and **F,** Two insulin bound (green ribbons) mouse INSR isoform A (PDB: 7STJ (asymmetrical conformation), and 7STH (symmetrical). The highlighted sections are rotated counterclockwise 90°. **G,** Saturated, four insulin bound (green ribbons) human INSR isoform A (PBD: 6PXV). The highlighted section is rotated counterclockwise 90°.

**Figure S7 (Related to Figure 4): Relationship among expression, binding and signaling scores** Boxplots of (**A**) cell surface expression, (**B**) insulin binding and (**C**) insulin signaling scores for previously functionally studied variants. The dashed horizontal line indicates the optimal cut off for binary classification of variants into wild-type and loss-of-function categories, determined through confusion matrix analysis. **D**, Venn diagram comparing results for categorical loss of expression, insulin binding, and signaling determined by applying cut-offs from A, B, and C. Analysis was limited to variants with scores generated for all three assays and an FDR < 0.05. This identified 2902, 4306, and 3782 loss-of-function variants for expression, insulin binding, and signaling, respectively. **E**. Classification of variant effects based on the Venn diagram results.

